# Distinct nanoscale dynamics of growth receptor complexes link hormone perception to rapid cell wall remodeling in *Arabidopsis thaliana*

**DOI:** 10.64898/2026.09.09.750143

**Authors:** Luiselotte Rausch, Aylin Balmes, Dominic Zoller, Jagmohan Singh, Ruairidh Macleod Davidson, Sven zur Oven-Krockhaus, Holger F. Bettinger, Tilman E. Schäffer, Klaus Harter

## Abstract

Cell elongation is a fundamental process allowing plants to change size and shape, a process governed by several growth promoting hormones. While hormones such as brassinosteroids (BRs) and phytosulfokines (PSKs) have been shown to play important roles in elongation growth, the early steps of signal perception remains elusive, especially with regard to PSK. Here we report a rapid mechanism by which PSK alter cell mechanical properties in elongating hypocotyls. Notably, this mode of action differs from that of BR. Making use of atomic force microscopy and fluorescence lifetime imaging microscopy we demonstrate how hallmarks of growing plant cells such as mechanical wall properties, porosity and apoplastic pH are differentially affected by PSK compared to BR. Using super-resolution microscopy, we show that the receptor complex components for BR and PSK display individual spatiotemporal movement and organization patterns. The BR receptor BRI1 transitions to a faster diffusive state upon ligand perception, while the PSK receptor PSKR1 associates in tighter clusters. The shared co-receptor BAK1 displays a selective decrease in cluster density after PSK treatment. Moreover, loss of functional BAK1 alters the nanoscale reorganization of PSKR1 after ligand perception. We found that the putative cell wall state sensor RLP44 is required for the observed changes, however the spatiotemporal dynamics of RLP44 are not altered during signaling. We propose a model of how cell walls are specifically tuned by BRI1- and PSKR1-centered signaling hubs as potential prerequisites for and during cell elongation initiation.

## Introduction

Plants have adapted intricate systems to adequately respond to their environment within short timeframes. The coordination of directed growth is fundamental to the acquisition of sunlight, water and nutrients. These processes are largely dependent on various phytohormones and require careful orchestration of changes in cell shape and mechanical cell properties.

One of the fastest elongating plant organs is the hypocotyl. After germination, hypocotyls expand rapidly through the soil in an acropetal manner (Gendreau et al., 1997; Krahmer et al., 2024). This process is effectively devoid of cell divisions and relies on extensive cell elongation (Gendreau et al., 1997). Therein, the epidermis is the tissue mainly restricting or allowing growth (Savaldi-Goldstein et al., 2007). On a cellular level, expansive cell growth results from the mechanical imbalance between internal turgor pressure and the surrounding wall. Turgor exerts an outward pressure, under which the wall yields by the rearrangement of microfibrils, resulting in irreversible extension and growth of the cell (Cosgrove, 2022; Ali et al., 2023). The rearrangement of cell wall components such as cellulose fibers is linked to wall-modifying enzymes such as expansins that are thought to cleave crosslinks between cellulose microfibrils and between cellulose and hemicellulose (McQueen-Mason et al., 1992; Samalova et al., 2022; Whitney et al., 2000). According to the ’acid growth theory’, cell wall-modifying enzymes are activated when the apoplastic pH is lowered by plant hormones such as auxin or brassinosteroids (Hager et al., 1971; Rayle et al., 1970; Großeholz et al., 2022). In more recent years, increasing evidence also suggests an involvement of pectins in acid growth processes (Arsuffi et al., 2018; Hocq et al., 2017), such as primordium formation in the shoot apical meristem (Braybrook et al., 2013).

Brassinosteroids (BRs) are plant steroid hormones that have been intensely investigated with regard to their role in cell elongation in organs such as the *Arabidopsis thaliana* (*A. thaliana*) root (Vukašinović et al., 2025; Lv et al., 2020). BRs are perceived by BRASSINOSTEROID-INSENSITIVE-1 (BRI1), a leucine-rich repeat receptor-like kinase (LRR-RLK) localized in the plasma membrane (PM) (Li et al., 1997; Belkhadir et al., 2006). Upon ligand binding, several BRI1-inhibiting proteins including BAK1-INTERACTING RECEPTOR-LIKE KINASE1 (BIR1) and BRI1 kinase inhibitor 1 (BKI1) are released, which allows interaction and transphosphorylation with the co-receptor BRI1-ASSOCIATED KINASE 1 (BAK1) (Bücherl et al., 2013; Bücherl et al., 2017; Li et al., 2002). Once the active receptor complex is established, two distinct pathways are initiated: The long-term transcriptional pathway involves translocation of the two master regulator transcription factors BRASSINAZOLE RESISTANT 1 (BZR1) and BR INSENSITIVE EMS49 SUPPRESSOR 1 (BES1) into the nucleus, which induce a multitude of gene-regulatory responses (He et al., 2002; Yin et al., 2002). Simultaneously, a rapid response takes place solely at the PM wherein the BRI1-BAK1 receptor complex upregulates the activity of the PM-resident proton pumps H^+^-ATP-ase 1 (AHA1) and H^+^-ATP-ase 2 (AHA2). This, in turn, leads to membrane hyperpolarization and extrusion of protons into the apoplast, lowering the apoplastic pH and thereby allowing for the onset of cell elongation (Caesar et al., 2011; Großeholz et al., 2022). This fast response takes place within 5-30 min in the meristematic and early elongation zone of *A. thaliana* roots (Großeholz et al., 2022).

The receptor complex for phytosulfokine (PSK) is structurally highly similar to the BRI1 receptor complex. The plant peptide PSK is perceived by the LRR-RLK PHYTOSULFOKINE RECEPTOR 1 (PSKR1) and PHYTOSULFOKINE RECEPTOR 2 (PSKR2) and similarly contributes to root growth and hypocotyl elongation (Stührwohldt et al., 2011; Kutschmar et al., 2009; Matsubayashi et al., 2006). Both PSKR1 and BRI1 possess a ligand-binding island domain, a single transmembrane domain and intracellular kinase activity (Hartmann et al., 2013; Kinoshita et al., 2005; Shinohara et al., 2007; Wang et al., 2008). Similarly to BRI1, PSKR1 interacts with BAK1, AHA1 and AHA2, however the two receptors do not interact with each other (Ladwig et al., 2015). This suggests that PSK-perception might be similar to BR-perception with regard to growth and cell elongation. While long-term effects of PSK have been studied (Matsubayashi et al., 2006; Stührwohldt et al., 2011; Kutschmar et al., 2009), a rapid cell physiological effect such as observed for BR-signaling has not been reported so far. Since both BRI1- and PSKR1-dependent signaling are promoted by their interaction with RECEPTOR-LIKE-PROTEIN 44 (RLP44), a putative cell wall sensor (Wolf et al., 2014; Holzwart et al., 2018; Holzwart et al., 2020), it seems likely that both pathways may relate to cell wall signaling. It is noteworthy that interaction between RLP44 and BRI1 is dependent on the phosphorylation state of RLP44, while RLP44 interacts with PSKR1 regardless of its phosphorylation state (Garnelo Gómez et al., 2021). Additionally, both receptors interact with members of the CYCLIC NUCLEOTIDE-GATED CHANNEL (CNGC)-family, namely CNGC10 and CNGC17 (Ladwig et al., 2015; Großeholz et al., 2022). These subtle differences may indicate specific differences between responses of these functionally intertwined receptor complexes. However, the question arises how specificity between BR- and PSK-signaling complexes is achieved and how these two signaling pathways are coordinated with regard to shared components and distinct cell physiological outputs. In recent years, the notion of specific organization and separation of signaling complexes within the PM has gained more interest as well as experimental support (Jaillais et al., 2020; Jaillais et al., 2024). Nowadays, the PM is seen as a heterogeneous space wherein specific proteins and lipids locally accumulate in specified signaling-hubs (Jaillais et al., 2024; Kusumi et al., 2005; Bücherl et al., 2017). Processes such as osmotic stress signaling and immune responses during viral infection were linked to PM-based mechanisms that rely on tight control of these signaling hubs, also referred to as nanodomains. (Smokvarska et al., 2023; Jolivet et al., 2025; Jaillais et al., 2024). In a recent study, the putative PM organizer HYPERSENSITIVE-INDUCED REACTION 2 (HIR2) has been shown to interact with LRR-RLKs such as BRI1 as well as playing an important role in the nanoscale organization of BAK1 in the PM (Weber et al., 2026). Therefore, it is highly probable that the strongly related receptor complexes of BRI1 and PSKR1 may be differently organized at nanoscale within the PM.

Since the PM is extremely densely populated with hundred millions of proteins per cell membrane (Heinemann et al., 2021), specialized super-resolution imaging methods are needed to properly characterize spatiotemporal behavior of individual PM components on the nanoscale. Single-particle tracking photoactivated localization microscopy (sptPALM) surpasses the diffraction limit, enabling single-molecule localization with a precision of approximately 20 nm (Betzig et al., 2006) and allows to determine the size of nanodomain-organized clusters as well as the diffusion coefficient and other mobility patterns of a chosen protein *in vivo* (zur Oven-Krockhaus et al., 2026; Rohr et al., 2024b). Therefore, sptPALM is a very promising approach for investigating how specific mechanisms during BR- and PSK-signaling are realized within the PM during early signaling events.

Here, we used atomic force microscopy (AFM) in combination with other biomechanical approaches to entangle how BR- and PSK-signaling affect cell physiological outputs in elongating hypocotyls on short timescales. AFM is a scanning probe method that relies on a flexible cantilever being deflected upon contact with the sample surface. The deflection can further be used to estimate mechanical properties of a material (Binnig et al., 1986; Colton et al., 1997; Rausch et al., 2026). In the context of plant cells the sample stiffness obtained by indentation can be linked to either cell turgor pressure of cell wall material properties, depending on indentation modality. We found that the mechanical properties of epidermal hypocotyl cells is differentially affected by the two pathways and investigated how changes in apoplastic pH and/or wall porosity may contribute to the biomechanical phenotypes observed in indentation experiments. To clarify how specificity between the closely related BRI1 and PSKR1 complexes is realized on nanodomain-level within the PM, we studied single-molecule mechanisms of relevant signaling complex components and discovered distinct spatiotemporal patterns of BRI1, PSKR1 and BAK1. We therefore propose a framework that links rapid cell elongation and cell wall remodeling to distinct PM-resident signaling domains, offering a new perspective on the regulation of cell elongation.

## Results

### PSK rapidly alters mechanical surface properties of elongating hypocotyls

Since PSKR1 interacts with the proton pumps AHA1 and AHA2 as well as the calcium channel CNGC17 and the putative cell wall state receptor RLP44, we hypothesized that PSK might contribute to elongation growth processes by rapidly altering mechanical properties of walls. We performed AFM indentations on periclinal cell walls of epidermal cells of 7-day-old etiolated hypocotyls of *A. thaliana* building on previously published protocols by Milani et al. (2011). We designed our measurements in shallow indentation depths where turgor becomes negligible (Vella et al., 2011; Vella et al., 2012) in order to ensure that we capture primarily changes in cell wall properties. We assume that the indentation stiffness extracted from shallow indentations on turgid cells to be mainly connected to the isotropic cell wall matrix (mostly pectins with secondary order contributions by fibrous wall polymer compression) (Rausch et al., 2026). Nevertheless, we adopted the term ’apparent stiffness’ for the material property resulting from data evaluation by the Hertz model to acknowledge the inherent complexity of AFM-based indentation (Rausch et al., 2026).

First, we tested whether our chosen settings capture the developmental differences of acropetal growth in etiolated hypocotyls by comparing the lower versus upper area of the hypocotyl (see Fig. 1A). In line with previous works by Seifert (2018), we found that the apparent stiffness at the upper part of the elongating hypocotyl was significantly higher compared to the lower part (Fig. 1A). To demonstrate that rapid physiological responses can be captured, we incubated hypocotyls with 5 nM fusicoccin (FC), a proton pump-activating fungal metabolite (Marre, 1979; de Boer, 2024), for 30 min and performed indentations in the central region of the hypocotyl. As predicted by the acid-growth theory, FC treatment leads to a reduction of measured apparent stiffness (Fig. 1B).

**Figure 1:**
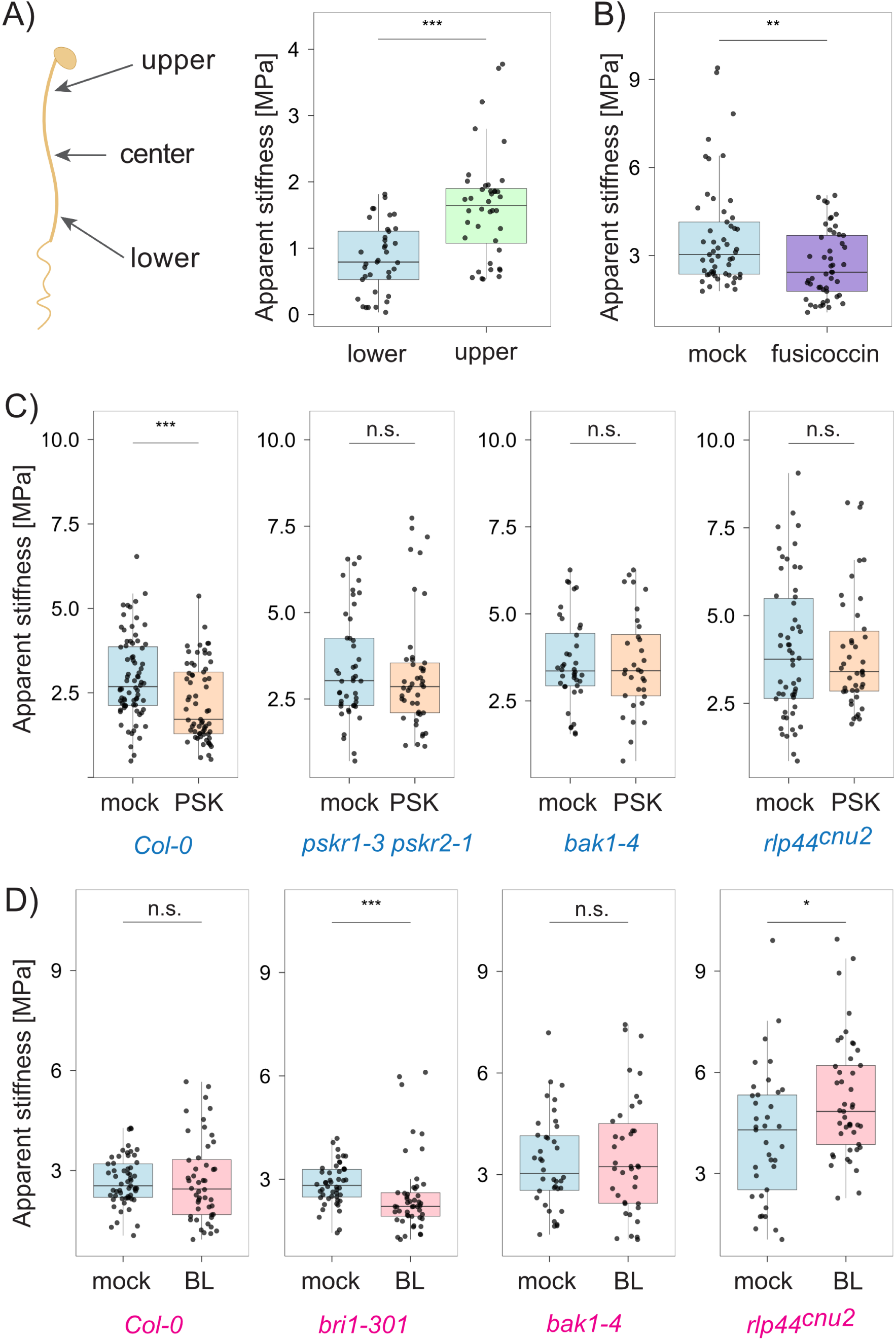
A) Schematic representation of 7-day-old etiolated *A. thaliana* seedlings used in atomic force microscopy experiments with relevant measurement positions indicated by arrows. Boxplot showing the apparent stiffness of epidermal hypocotyl cells of etiolated seedlings measured at the bottom versus top position. Each data point represents the median apparent stiffness of a single epidermal cell. Data pooled from 3 measurement days. B) Boxplot showing the apparent stiffness of epidermal hypocotyl cells of etiolated seedlings measured at the central position after 30 min treatment with 5 nM Fusicoccin or mock treatment. Each datapoint represents the median apparent stiffness of a single epidermal cell. Data pooled from 5 measurement days. C) Boxplot showing the apparent stiffness of epidermal hypocotyl cells of etiolated seedlings measured at the central position after 30 min treatment with 100 nM phytosulfokine (PSK) or mock treatment in wildtype and relevant signaling mutants (from left to right: *Col-0*, *pskr1-3 pskr2-1*, *bak1-4*, *rlp44^cnu2^*). Mock shown in light blue and PSK shown in light orange for all genotypes. Each datapoint represents the median apparent stiffness of a single epidermal cell. Data pooled from 6 measurement days for *Col-0* ; 5 measurement days pooled for all other genotypes. D) Boxplot showing the apparent stiffness of epidermal hypocotyl cells of etiolated seedlings measured at the central position after 30 min treatment with 10 nM brassinolide (BL) or mock treatment in wildtype and relevant signaling mutants (from left to right: *Col-0*, *bri1-301*, *bak1-4*, *rlp44^cnu2^*). Mock shown in light blue and BL shown in light pink for all genotypes. Each datapoint represents the median apparent stiffness of a single epidermal cell. Data pooled from 5 measurement days. All statistical analyses were performed using custom-made R scripts and the Wilcoxon test distribution (applicability checked prior using a Levene’s test and a Shapiro-Wilk test). n *≥* 32 for each condition. p *≤* 0.001 (***); p *≤* 0.01 (**); p *≤* 0.05 (*); p *>* 0.05 (n.s.). Abbreviations: Phytosulfokine (PSK), brassinolide (BL).

Next, we treated hypocotyls with 100 nM PSK for 30 min and found a strong decrease in apparent stiffness in the central region of the hypocotyl (Fig. 1C). Importantly, this response is not taking place in relevant signaling mutant lines such as the T-DNA insertion lines *pskr1-3 pskr2-1* and *bak1-4* (Fig. 1C). The mutant line *rlp44^cnu2^*, in which RLP44 is non-functional due to a premature stop-codon (Wolf et al., 2014), was also non-responsive to PSK (Fig. 1C). Therefore, all these receptor components are contributing to this rapid PSK-dependent cell wall-related phenotype.

We performed similar indentations after treatment with 10 nM Brassinolide (BL) for 30 min. We did not observe significant changes in apparent stiffness in wildtype plants (Fig. 1D). The mutant line *bri1-301*, in which BRI1’s kinase activity is reduced (Lv et al., 2020; Zhang et al., 2018), was hypersensitive to BL treatment with respect to apparent cell wall stiffness. BL treatment showed no effect in *bak1-4*, and an increase in apparent stiffness in *rlp44^cnu2^* (Fig. 1D). Since these measurements were all conducted in the central region of the hypocotyl, we speculated that BL-signaling may occur in a position-dependent manner, which may explain the lack of an effect on apparent stiffness in our measurements conducted in the central region. We therefore performed indentations after BL-treatment separately in the top and bottom regions of the hypocotyl; however, these experiments remained inconclusive (data not shown).

### PSK and BL induce rapid cell wall acidification

In order to investigate the effects observed by AFM in more detail, we used CarboTags (Besten et al., 2025) for fluorescence lifetime imaging microscopy (FLIM) experiments in the cell walls of the *A. thaliana* hypocotyl. CarboTag (CT) is a modular tool developed for cell wall imaging that consists of a pyridinium boronic acid targeting motif containing an alkyne group that can be clicked with azide-functional fluorophores. The fluorescence lifetime (FLT) of the green fluorescent dye Oregon Green (OG) has been previously shown to be a suitable readout for pH (Barnoy et al., 2019; Besten et al., 2025). Based on the synthesis protocols published in Besten et al. (2025), we synthesized CT linked to OG for measuring changes in cell wall pH (see Materials and Methods). We conducted all CT-OG experiments in a two-step setup: 7-day-old etiolated seedlings were incubated in the CT-OG staining solution for 1 h, followed by transfer into the treatment solution for 30 min before performing the FLIM measurement (Fig. 2A). This two-step incubation was chosen to accomodate both the optimum staining duration of the CT (1 h) and the short incubation required to address early, non-transcriptional signaling events, as studied in our AFM experiments. Fluorescence was exclusively detected at the cell wall, indicating successful staining (Fig. 2B).

**Figure 2:**
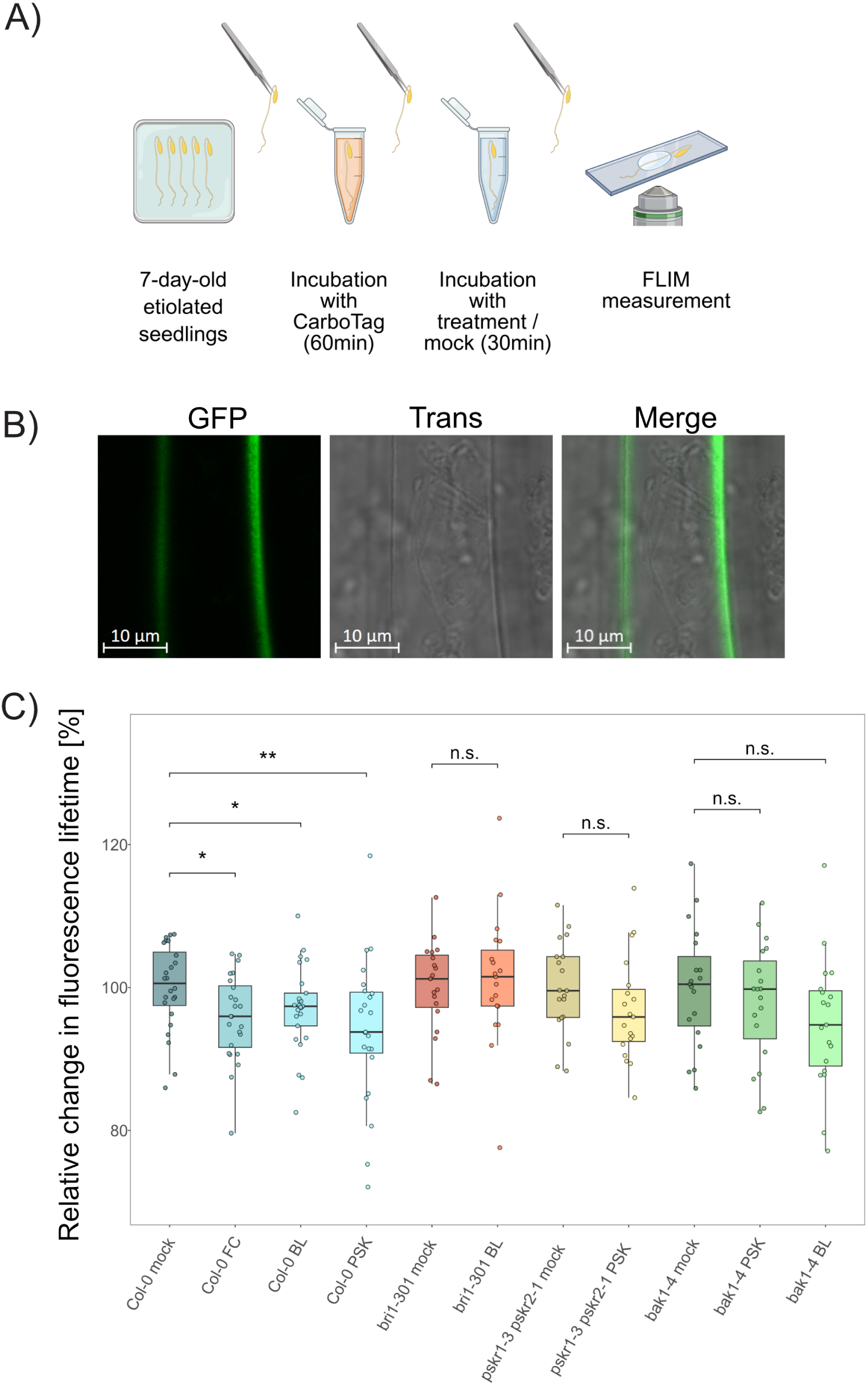
A) Schematic workflow of fluorescence lifetime imaging microscopy (FLIM) measurements performed on CarboTag (CT)-stained etiolated hypocotyls. Created in BioRender. Rausch, L. (2026) https://BioRender.com/f33tz3a. 7-day-old etiolated seedlings were carefully transferred with tweezers into a 10 µM CT staining solution for 60 min, followed by incubation in fusicoccin (FC), brassinolide (BL), phytosulfokine (PSK) or mock treatment for 30 min before FLIM imaging. B) Exemplary confocal image of CarboTag-Oregon Green (CT-OG)-stained hypocotyl cells in *A. thaliana* showing cell wall-localized staining by the CT-OG dye. Samples prepared as described above. Left = GFP channel, middle = transmitted light channel, right = overlay of GFP and transmission channels. Scalebar = 10µm. C) Boxplot showing the relative change in cell wall pH of CT-OG stained etiolated hypocotyls after FC, BL and PSK treatment in wildtype plants and relevant signaling mutants. The data derived from the mock treatments of the respective line were set to 100. Each datapoint represents the relative fluorescence lifetime derived from individual seedlings by scanning an area of 35.42 µm x 35.42 µm and selecting the apoplastic regions using the free ROI tool in SymPhoTime 64 (Pico-Quant GmbH, Berlin, Germany). Data pooled from 4 measurement days; all conditions were measured in *≥*3 samples on each measurement day. All statistical analyses were performed using custom-made R scripts and the Wilcoxon test distribution (applicability checked prior using a Levene’s test and a Shapiro-Wilk test). n *≥* 19 for each condition. p *≤* 0.001 (***); p *≤* 0.01 (**); p *≤* 0.05 (*); p *>* 0.05 (n.s.). Abbreviations: fluorescence lifetime imaging microscopy (FLIM), CarboTag (CT), CarboTag-Oregon Green (CT-OG), fusicoccin (FC), brassinolide (BL), phytosulfokine (PSK).

First, we tested the effect of 30 min incubation in 5 nM FC and found the expected decrease in wall pH (see Fig. 2C), in line with the results published in Besten et al. (2025). We found a similar drop in wall pH after BL application that aligns with previous works (Caesar et al., 2011; Großeholz et al., 2022), and an even stronger decrease after PSK treatment (Fig. 2C). When treating the *bri1-301* and *bak1-4* mutants with BL, no significant effect was observed (although there was still some residual effect in *bak1-4*, p = 0.064). These results demonstrate that BL-induced lowering in pH is functional in the same region, where AFM indentations were performed. PSK-induced acidification of the cell wall was also absent in the *pskr1-3 pskr2-1* and *bak1-4* mutants. Therefore both BL and PSK induce rapid cell wall pH decrease in etiolated hypocotyls via their respective receptor complexes. Since both BRI1 and PSKR1 interact with the proton pumps AHA1 and AHA2, a mode of action that aligns with acid growth theory appears very likely. However, it raises the question why FC and PSK lead to a clear phenotype in the apparent wall stiffness, but BL does not.

### PSK rapidly alters cell wall porosity

Acid growth theory primarily addresses mechanisms linked to cellulose reorganization by pH-responsive enzymes like expansins (Cosgrove, 2024; Du et al., 2020). However, the wall also consists of other components such as pectins and xyloglucan (Cosgrove, 2022). It was recently suggested that the link between cellular pH and cellular growth as proposed by acid growth theory may be less straight-forward than previously thought (Krupař et al., 2026). In addition, contributions of pectin to acid-driven growth processes have been observed (Hocq et al., 2017; Braybrook et al., 2013). To explore different wall properties that may contribute to the phenotypes observed in AFM, we investigated how so-called cell wall porosity (referring to the mesh size of the pectin network; Besten et al. (2025)) changes under our treatments. The molecular rotor phenyl-BODIPY (boron dipyrromethene, BDP) is a green fluorescent probe that can monitor structural properties in its environment via its FLT (Michels et al., 2020; Michels et al., 2022; Liu et al., 2020). When intramolecular rotation of the molecular rotor is restricted by its local environment, access to non-radiative decay pathways is reduced, leading to an increase in FLT. On the other hand, FLT of BDP is low when the surroundings allow free rotation. We coupled BDP to the CT backbone via click chemistry and performed stepwise incubation similar to CT-OG experiments described above (1 h in the CT-BDP solution, followed by 30 min incubation in BL, PSK or control treatment). Fluorescence was exclusively detected at the cell wall (Fig. 3A). To verify the performance of our dyes in FLIM measurements, we included a control experiment based on Besten et al. (2025). Specifically, we compared seedlings grown on 3 nM isoxaben (ISX), a cellulose synthesase inhibitor, to seedlings grown on regular medium. We confirmed the results of Besten et al. (2025), showing that ISX treatment increases the FLT of CT-BDP, reflecting higher cell wall porosity (see Fig. 3B).

**Figure 3:**
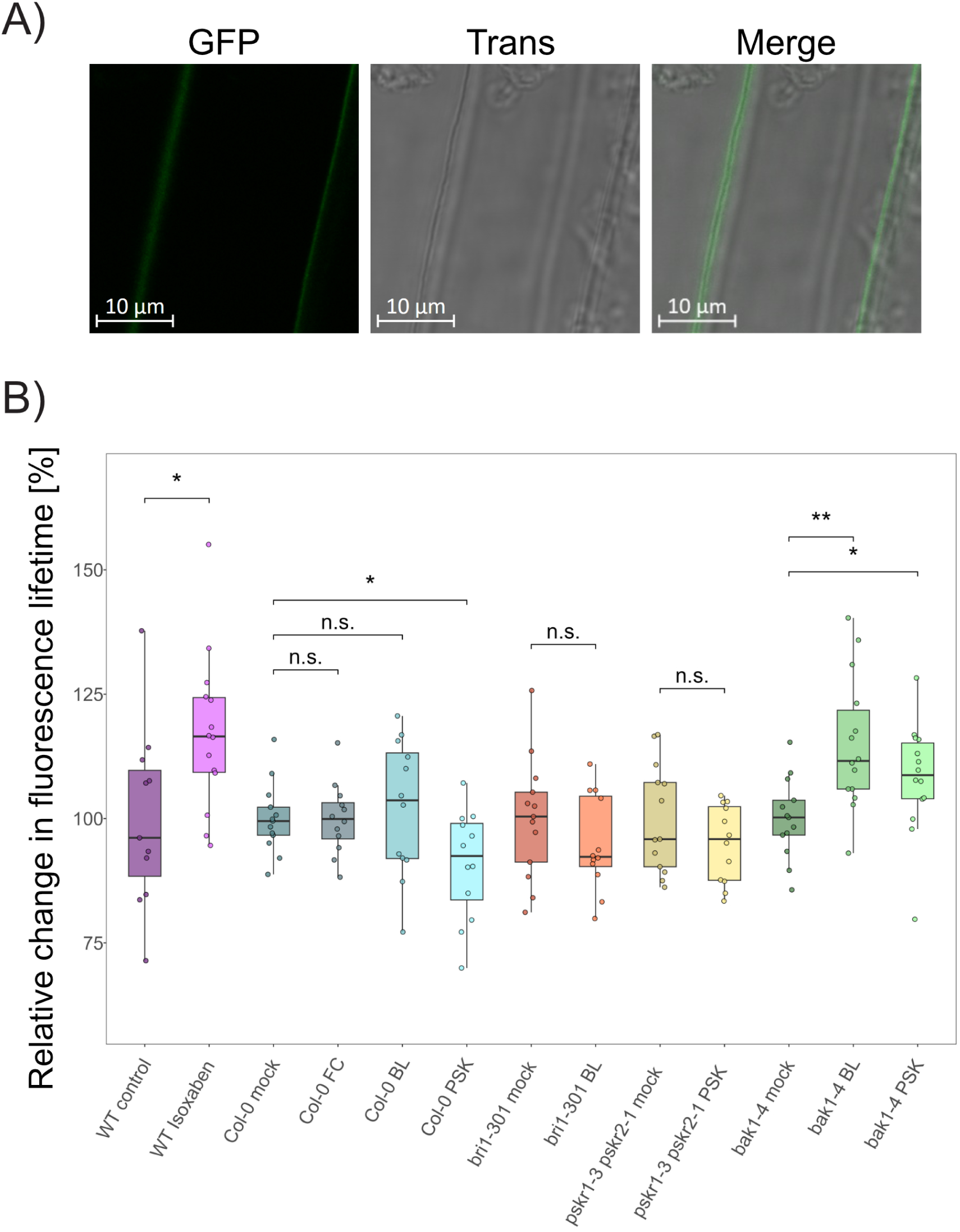
A) Exemplary confocal image of CarboTag-BODIPY (CT-BDP)-stained hypocotyl cells in *A. thaliana* showing cell wall-localized staining by the CT-BDP dye. Samples prepared as described in the figure legend for Fig. 2A, using CT-BDP at 10 µM for staining. Left = GFP channel, middle = transmitted light channel, right = overlay of GFP and transmission channels. Scalebar = 10 µm. B) Boxplot showing the relative change in cell wall porosity of CT-BDP stained etiolated hypocotyls after fusicoccin (FC), brassinolide (BL) or phytosulfokine (PSK) treatment in wildtype (WT) plants and relevant signaling mutants. The data derived from the mock treatments of the respective line were set to 100. Each datapoint represents the relative fluorescence lifetime derived from individual seedlings by scanning an area of 35.42 µm x 35.42 µm and selecting the apoplastic regions using the free ROI tool in SymPhoTime 64 (PicoQuant GmbH, Berlin, Germany). Data pooled from 3 measurement days; all conditions were measured in *≥*3 samples on each measurement day. All statistical analyses were performed using custom-made R scripts and the Wilcoxon test distribution (applicability checked prior using a Levene’s test and a Shapiro-Wilk test). n *≥* 11 for each condition. p *≤* 0.001 (***); p *≤* 0.01 (**); p *≤* 0.05 (*); p *>* 0.05 (n.s.). Abbreviations: CarboTag-BODIPY (CT-BDP), fusicoccin (FC), brassinolide (BL), phytosulfokine (PSK), wildtype (WT).

Next, we tested the short-term effect of 100 nM PSK and 10 nM BL on cell wall porosity in etiolated seedlings. To clarify whether general proton pump activation and drop in pH as observed after FC is related to cell wall porosity, we included FC treatment into our CT-BDP experiments. Only PSK treatment led to a decrease in FLT (ergo an increase in mesh size/porosity, Fig. 3B). BL and FC treatment did not alter cell wall porosity, but BL treatment lead to a slight spread towards higher FLTs. The mutant lines *bri1-301* and *pskr1-3 pskr2-1* did not react to treatment with respective ligands. Interestingly, *bak1-4* showed an increase in FLT to both BL and PSK, with a stronger effect after BL application. We could therefore show that only PSK signaling has a phenotype related to cell wall porosity. Therefore, PSK perception probably differs from BR signaling in terms of pectin network alteration.

### PSKR1 and BAK1 reorganize within nanodomain structures upon PSK treatment

All signaling components that were investigated by AFM and CT FLIM measurements are PM-localized. The concept that signaling specificity between closely related PM-resident processes may be realized by spatiotemporal segregation within the PM prompted us to investigate how BL- and PSK-signaling components behave immediately after ligand perception. To access nanoscale parameters of individual PM-proteins, we used sptPALM to characterize the behavior of individual proteins involved in early BR- and PSK-signaling on the single-molecule level. We expressed BRI1, PSKR1, and RLP44 fused to the photoconvertible fluorophore mEos3.2 under their native promotors in *A. thaliana* in their respective mutant backgrounds (except for RLP44 that was expressed in *Col-0*). Exemplary images of the four target proteins during sptPALM tracking in the central region of 7-day-old etiolated hypocotyls of *A. thaliana* in epidermal cells are depicted in Fig. 4A. We first investigated how PSKR1, BAK1 and RLP44 organize within the PM and how their behavior changes upon PSK treatment. Analysis of the raw data was performed using the software package OneFlowTraX (Rohr et al., 2024b) with regard to protein diffusion coefficient, cluster size and distribution of subpopulations. We found that all three proteins show a Gaussian distribution of diffusion coefficients (See Suppl. Fig. S6A-C) and no alteration of their lateral mobility upon PSK treatment (Fig. 4B). We evaluated the area of PSKR1-, BAK1- and RLP44-clusters via Voronoi tessellation (Levet et al., 2025). Cluster sizes were distributed log-normally as observed in similar works (Rohr et al., 2024a). While the absolute size of clusters did not change after PSK treatment (Suppl. Fig. S1A-C), we found that the density of tracked particles within clusters was significantly altered (the number of tracks within each area classified as a cluster is directly provided by OneFlowTraX software package): the number of tracks per µm^2^ in PSKR1 clusters significantly increased after PSK treatment (log-transformed distributions shown for ease in Fig. 4C; raw data distributions for PSKR1, BAK1 and RLP44 shown in Suppl. Fig. S2A-C). Meanwhile, cluster density of BAK1 was decreased upon PSK application, and RLP44 cluster density remained unaltered. Since PSKR1 and BAK1 interaction is an integral part of PSK-sensing (Ladwig et al., 2015), the assumption of them interacting in pre-formed nanodomains appears plausible. We also found the number of clusters formed by BAK1 to be drastically lower than the number of PSKR1-clusters, which is in line with the recent work by von Arx et al. (2026), who found a similarly sparse clustering behavior and proposed a model in which BAK1 is recruited into LRR-RLK clusters such as FLAGELLIN-SENSING 2 in a BIR3-dependent manner.

**Figure 4:**
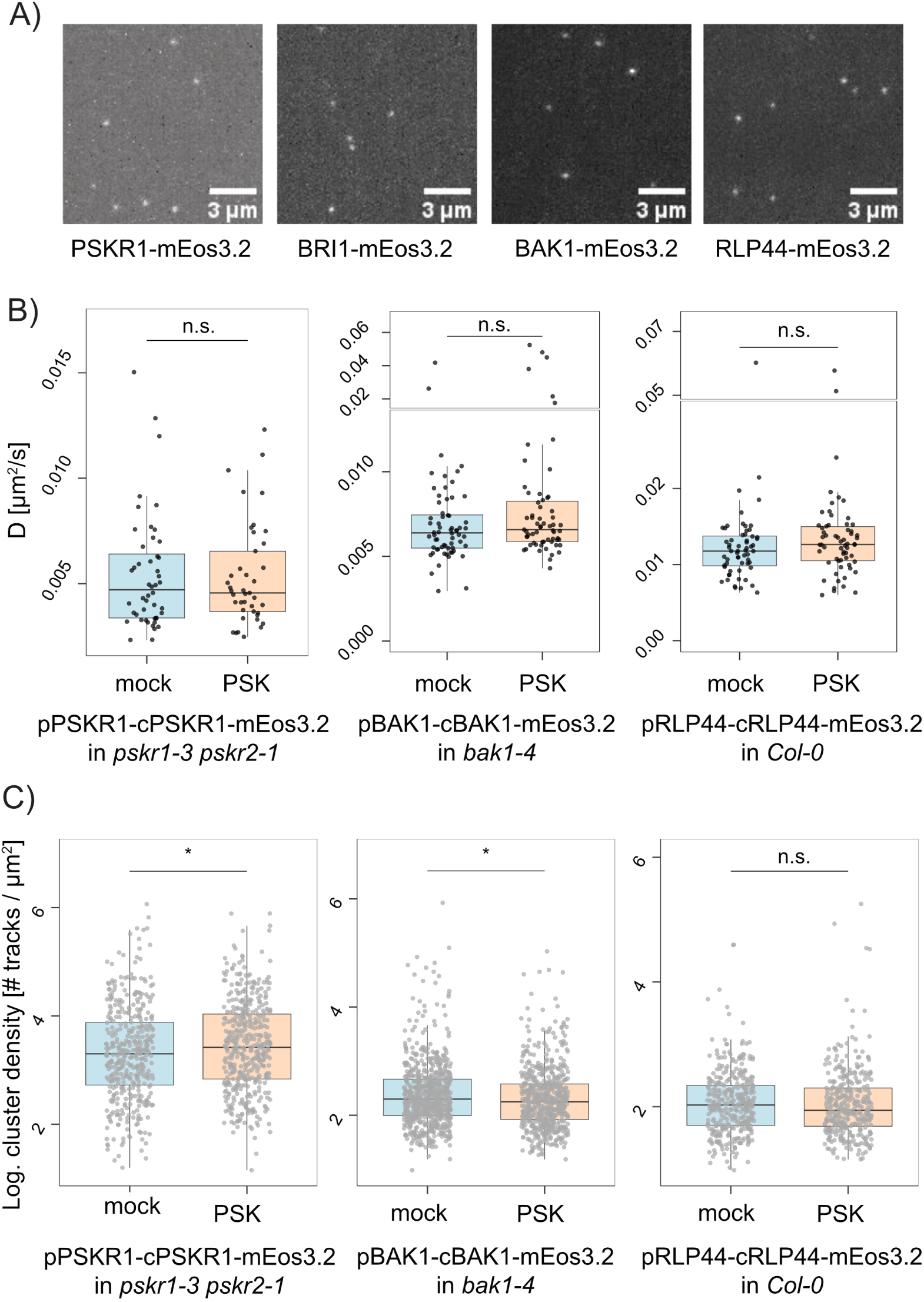
Nanoscale spatiotemporal behavior of phytosulfokine (PSK) and brassinosteroid (BR) signaling components in epidermal cells of 7-day-old etiolated hypocotyls in *A. thaliana* A) Representative images of PSKR1-mEos3.2, BRI1-mEos3.2, BAK1-mEos3.2 and RLP44-mEos3.2 during single-particle tracking photoactivated localization microscopy imaging in epidermal hypocotyl cells of *A. thaliana*. Movies were recorded in a 12.8 µm x 12.8 µm field of view. B) Diffusion coefficients of PSKR1-mEos3.2, BAK1-mEos3.2 and RLP44-mEos3.2 after 30 min incubation with 100 nM PSK or control treatment in epidermal hypocotyl cells of etiolated *A. thaliana* seedlings. Each datapoint represents the peak diffusion coefficient (D) of a single cell, calculated from a normal fit of the diffusion coefficient distribution within each sample. The control data shown here for BAK1 and RLP44 are shared with the according figures in figure 5A, as both experiments were performed concurrently under identical conditions. C) Cluster density (log-transformed) of PSKR1-mEos3.2, BAK1-mEos3.2, RLP44-mEos3.2 after 30 min incubation with 100 nM PSK or control treatment in epidermal cells of etiolated hypocotyls in *A. thaliana*. Their respective cluster areas were calculated via Voronoi tessellation (Levet et al., 2025). Density was determined as the number of tracks within a cluster divided by its area and then log-transformed in custom-made analysis pipelines in RStudio. Diffusion and cluster data shown in Fig. The control data shown here for BAK1 and RLP44 are shared with the according figures in figure 5B, as both experiments were performed concurrently under identical conditions. 4B-C are pooled from 4 measurement days; all conditions were measured in *≥*3 samples per condition on each measurement day. The number of cells per condition in which D and log-transformed cluster densities were evaluated is n *≥* 41 for PSKR1, n *≥* 63 for BAK1 and n *≥* 60 for RLP44. All statistical analyses were performed using custom-made R scripts and the Wilcoxon test distribution (applicability checked prior using a Levene’s test and a Shapiro-Wilk test). p *≤* 0.001 (***); p *≤* 0.01 (**); p *≤* 0.05 (*); p *>* 0.05 (n.s.). Abbreviations: Phytosulfokine (PSK), brassinosteroid (BR), peak diffusion coefficient (D).

To further characterize motion patterns of PSK-signaling related proteins, we employed the divide-and-conquer moment scaling spectrum (DC-MSS) algorithm established by Vega et al. (2018) that classifies motion trajectories into directed diffusion, free diffusion, confined diffusion or immobility, as previously described in Rohr et al., 2024a. Our results show that the predominantly confined diffusion behavior of PSKR1 (*≈* 51 %) is not altered by PSK treatment (Suppl. Fig. S8B). BAK1, on the other hand, shows *≈* 50 % free diffusion and *≈* 42 % confined diffusion (Suppl. Fig. S8C). Upon PSK treatment, BAK1 behavior showed a tendency towards increased free diffusion and less confined motion (Suppl. Fig. S8C). Lastly, RLP44 showed the highest percentage of free diffusion (*≈* 71 %, Suppl. Fig. S8D), which only marginally increased upon PSK treatment. Therefore, the LRR-RLK PSKR1, its co-receptor BAK1 and the receptor-like protein RLP44 all display individual motion patterns, but only BAK1’s relative motion patterns are affected by PSK.

### BRI1 switches into faster population upon BL treatment

To test whether BR perception differs from PSK-perception with regard to PM-localized spatiotemporal patterns, we analyzed the behavior of BRI1, BAK1 and RLP44 during signal initiation. After 30 min incubation in 10 nM BL, we found that only BRI1 showed altered diffusion, with increased lateral mobility (Fig. 5A). Intruigingly, when assessing the distribution of diffusion coefficients within the raw data, we noticed that the BRI1 datapool consisted of two diffusion populations. When exposed to BL, BRI1’s faster population increases (Fig. 5C). It is important to note that we observed two populations in each individual tracking movie (Suppl. Fig. S5A). Therefore the increase of the faster population is not an artifact caused by a subset of cells when pooling the data. We also quantified the relative amount of tracks residing within the faster and the slower population using the component-fit option of OneFlowTraX (Rohr et al., 2024b). Overall, the fraction of tracks within the slower population significantly decreased upon BL treatment (Suppl. Fig. S5B). We observed this specific effect only for BRI1; BAK1 and RLP44 displayed a single population (Suppl. Fig. S5C-D) that did not alter its diffusion coefficient (Fig. 5A).

**Figure 5:**
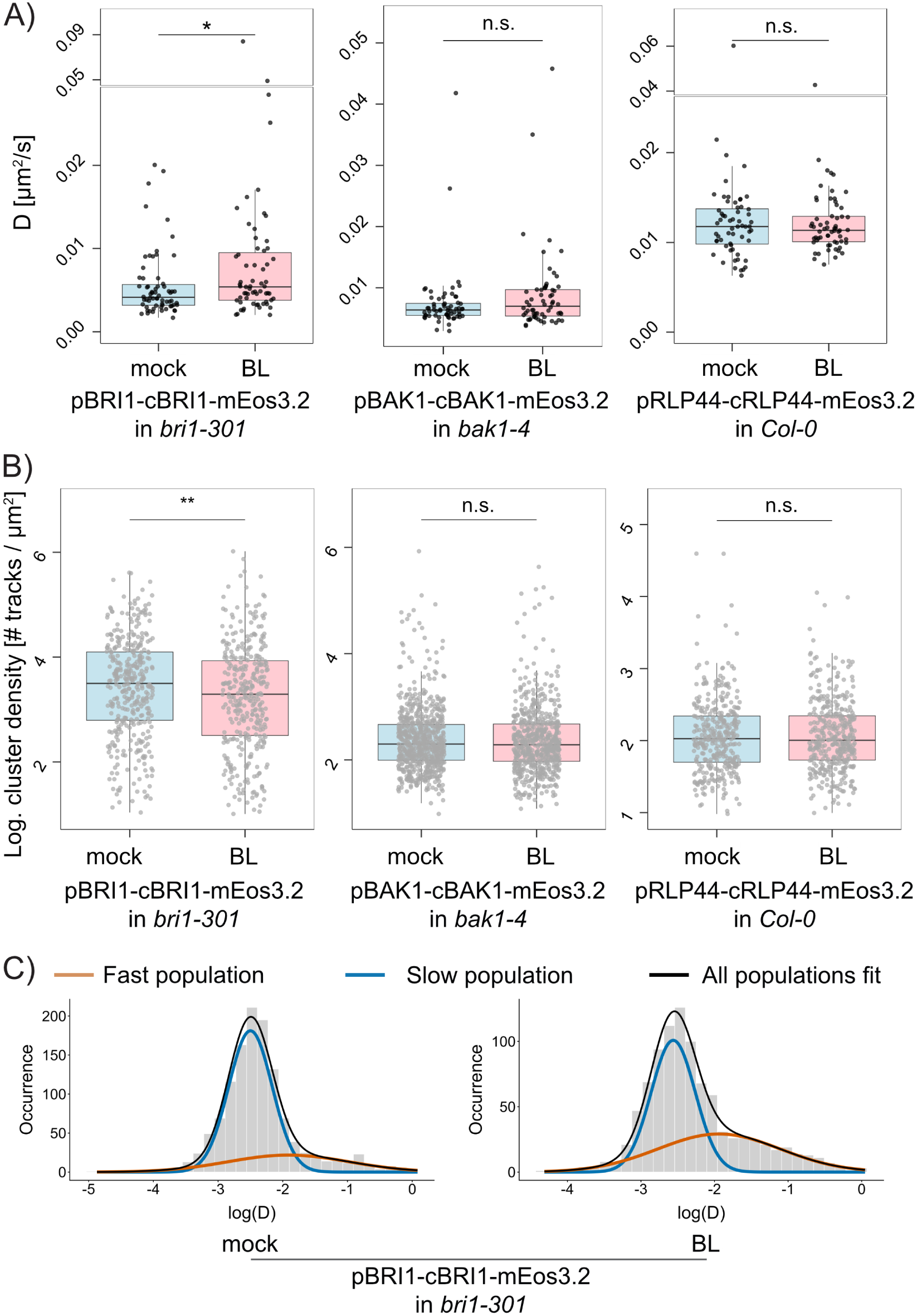
Nanoscale spatiotemporal behavior of brassinosteroid (BR) signaling components in epidermal cells of etiolated hypocotyls in *A. thaliana* A) Diffusion data for BRI1-mEos3.2, BAK1-mEos3.2 and RLP44-mEos3.2 after 30 min incubation with 10 nM brassinolide (BL) or control treatment in epidermal hypocotyl cells of etiolated *A. thaliana* seedlings. Each datapoint represents the peak diffusion coefficient (D) of a single cell, calculated from a normal fit of the diffusion coefficient distribution within each sample. Control data for BAK1 and RLP44 are identical to those in figure 4B due to simultaneous execution under identical conditions. B) Cluster density (log-transformed) of BRI1, BAK1, RLP44 in epidermal cells of etiolated hypocotyls after 30 min incubation with 10 nM BL or control treatment in epidermal cells of etiolated hypocotyls in *A. thaliana*. Cluster area was calculated via Voronoi tessellation (Levet et al., 2025). Density was determined as the number of tracks within a cluster divided by its area and then log-transformed in custom-made analysis pipelines in RStudio. Control data for BAK1 and RLP44 are identical to those in figure 4C due to simultaneous execution under identical conditions. C) Diffusion coefficient distribution of BRI1 after control (left) and BL (right) treatment. The two populations are indicated by the colored curves obtained via a two-component Gaussian mixture model. Both histograms were obtained from the pooled data for each condition on one exemplary measurement day (with n *≥* 19 cells per condition). Diffusion and cluster data shown in Fig. 5A-B are pooled from 4 measurement days; all conditions were measured in *≥*3 samples per condition on each measurement day. The number of cells per condition in which D and log-transformed cluster densities were evaluated is n *≥* 63 for BRI1, n *≥* 62 for BAK1 and n *≥* 60 for RLP44. All statistical analyses were performed using custom-made R scripts and the Wilcoxon test distribution (applicability checked prior using a Levene’s test and a Shapiro-Wilk test). p *≤* 0.001 (***); p *≤* 0.01 (**); p *≤* 0.05 (*); p *>* 0.05 (n.s.). Abbreviations: Brassinosteroid (BR), brassinolide (BL), peak diffusion coefficient (D).

As for the investigation of PSK, the distribution of cluster sizes followed a log-normal distribution for BRI1, BAK1 and RLP44 (Suppl. Fig. S3A-C). Upon BL treatment, track density of BRI1 within BRI1 clusters was significantly lower (log-transformed data shown in Fig. 5B, raw cluster density data shown in Suppl. Fig. S4A). Meanwhile, no such behavior was observed for BAK1 and RLP44, where cluster sizes and density remained unaltered by BL treatment (Fig. 5B, Suppl. Fig. S4B-C). As in our previous sptPALM experiments on PSK-perception, the number of clusters was higher for BRI1 compared to BAK1 and RLP44. The putative cell wall sensor RLP44 displayed a diffusion coefficient significantly higher than both BAK1 and the LRR-RLKs BRI1 and PSKR1 (Fig. 4B and Fig. 5A). We analyzed the motion patterns of BRI1, BAK1 and RLP44 during BR-signaling and found that BRI1 displays mostly confined behavior, similar to PSKR1 (Suppl. Fig. S8A-B). Upon BL treatment, we found a shift towards even more confined diffusion for BRI1, which highlights that decreased clustering is not automatically linked to more free diffusion. BAK1 and RLP44 showed no change in their motion behavior upon BL treatment (Suppl. Fig. S8C-D). Since BAK1 is required for the receptor complex assembly and signal initiation in both BR and PSK signaling (Caesar et al., 2011; Ladwig et al., 2015; Wang et al., 2008; Wang et al., 2015a), as confirmed by the AFM and CT-FLIM experiments in this work, it is noteworthy that only PSK triggered a change in spatiotemporal patterns of BAK1. RLP44 is known to interact with BRI1 in a phospho-state-dependent manner, therefore it will be of special interest to determine whether these two interactions are reflected in specific single-molecule patterns, which may be a subject for future dual color sptPALM studies.

### BAK1 is required for nanoscale organization of PSKR1 during early signaling events

Our sptPALM experiments revealed that PSK and BR perception components display individual nanoscale organization and mobility patterns. While these results demonstrate that specific physiological outputs are accompagnied by distinct mechanisms of PM-resident proteins, the causal necessity of these nanoscale processes for successful signaling is not fully established. To functionally link our single-molecule observations to the observed cellular output, we performed the same sptPALM experiment on PSKR1-mEos3.2 after PSK-treatment but in the *bak1-4* mutant background as a representative example. In the absence of the co-receptor BAK1, the receptor PSKR1 displayed altered nanoscale behavior. The general distributions of cluster sizes and diffusion coefficients followed log-normal and Gaussian distributions respectively, as observed in the previous measurements (Suppl. Fig. S7A-B). However, we observed a strong increase in lateral mobility of PSKR1 after ligand treatment that was not observed before (Fig. 6A). Moreover, the increase in PSKR1 cluster density that was found in WT-like conditions (Fig. 4C) was strongly altered in the *bak1-4* mutant background (Fig. 6B, Suppl. Fig. S7C). The motion type analysis revealed that PSKR1 showed roughly 10 % increase in immobility in absence of the ligand (Suppl. Fig. S7D) compared to the WT-like conditions described before (Suppl. Fig. S8B). Meanwhile, confined and free diffusion were reduced (Suppl Fig. S7D and S8B). These results show that in the absence of BAK1, where PSK-induced decrease of apparent stiffness and alteration of cell wall pH and porosity are hindered, the PM-resident single-molecule processes of PSKR1 are obstructed. In this situation, ligand treatment triggered a mobilization of the PSKR1 receptor and cluster density decrease, which may indicate that BAK1 acts as a molecular glue that is required for successful clustering and thereby enables effective signaling.

**Figure 6:**
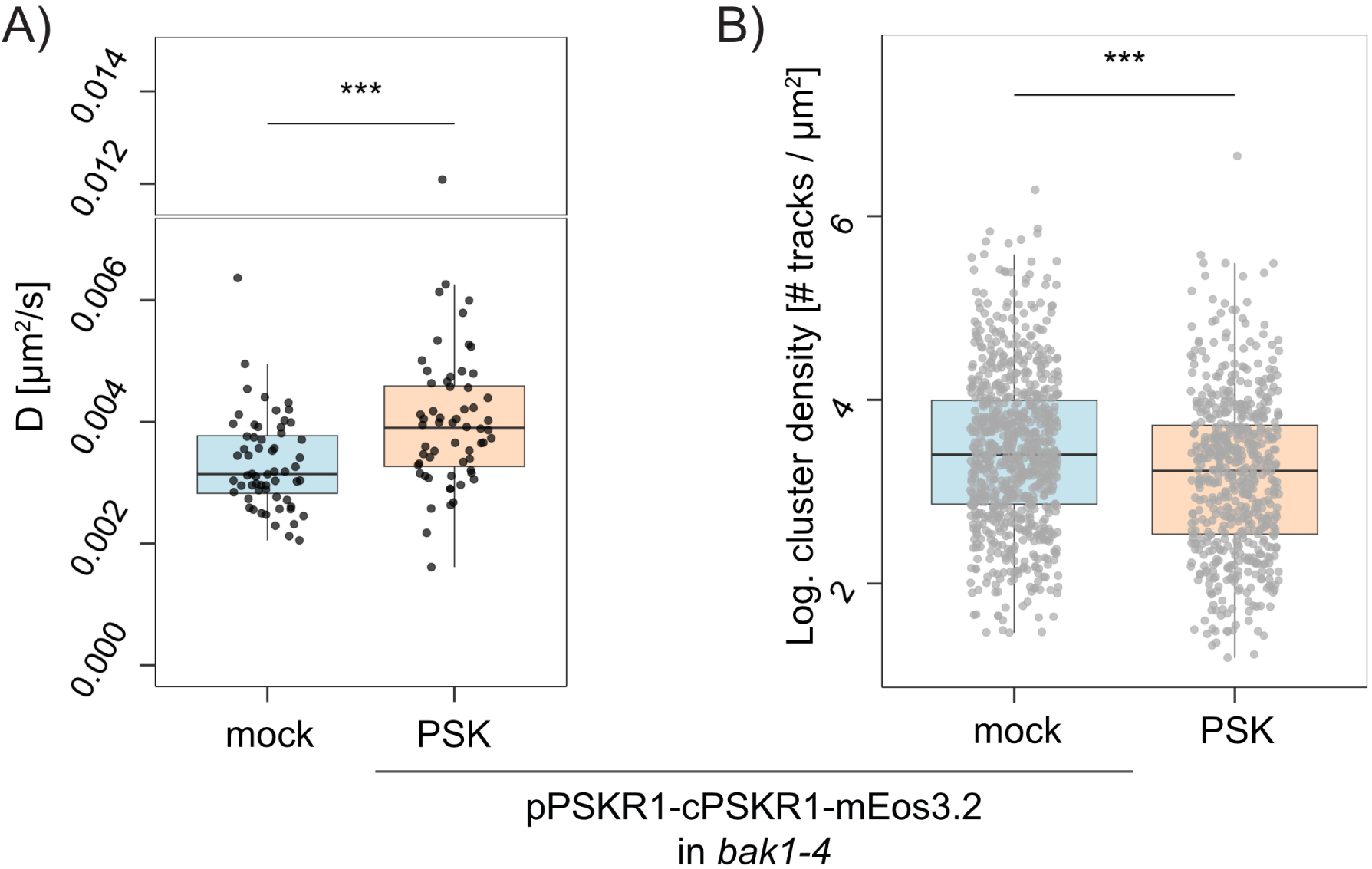
Nanoscale spatiotemporal behavior of PSKR1 during Phytosulfokine (PSK) treatment in epidermal cells of 7-day-old etiolated hypocotyls in the mutant background *bak1-4*. A) Diffusion coefficients of PSKR1-mEos3.2 after 30 min incubation with 100 nM PSK or control treatment in epidermal hypocotyl cells of etiolated *A. thaliana* seedlings (pPSKR1-cPSKR1-mEos3.2 in *bak1-4*). Each datapoint represents the peak diffusion coefficient (D) of a single cell, calculated from a normal fit of the diffusion coefficient distribution within each sample. B) Cluster density (log-transformed) of PSKR1-mEos3.2 after 30 min incubation with 100 nM PSK or control treatment in epidermal cells of etiolated hypocotyls in *A. thaliana* (pPSKR1-cPSKR1-mEos3.2 in *bak1-4*). Their respective cluster areas were calculated via Voronoi tessellation (Levet et al., 2025). Density was determined as the number of tracks within a cluster divided by its area and then log-transformed in custom-made analysis pipelines in RStudio. All data is pooled from 4 independent measurement days. The number of cells per condition in which D and log-transformed cluster densities were evaluated is n = 60. All statistical analyses were performed using custom-made R scripts and the Wilcoxon test distribution (applicability checked prior using a Levene’s test and a Shapiro-Wilk test). p *≤* 0.001 (***); p *≤* 0.01 (**); p *≤* 0.05 (*); p *>* 0.05 (n.s.). Abbreviations: Phytosulfokine (PSK).

### Proposed model of rapid BL and PSK signaling specificity

Here we studied how two closely related pathways involved in cell elongation in etiolated hypocotyls affect rapid physiological responses in the plant cell wall and how their signaling complex components behave during signal initiation on a single-molecule level in the PM. When summarizing all data obtained from AFM indentations, CT-based FLIM measurements and sptPALM measurements, we obtained distinct behaviors of the two closely related complexes for BR and PSK perception. We therefore propose the following model of how these two specific reaction pathways are carried out during ligand perception and signal initiation.

Our AFM measurements demonstrate that PSK triggers rapid alteration of the cell wall, a process that depends on the receptor complex components PSKR1, BAK1 and RLP44 (Fig. 1C). By using CT-FLIM, we found that this process probably relies on lowering the cell wall pH as well as changes in the pectin network (Fig. 2C, 3B). When investigating how these processes may be coordinated on the nanoscale level, we observed that the receptor PSKR1 exhibits primarily confined diffusion (Suppl. Fig. S8B) and is organized in nanodomains that show increased receptor track density upon ligand treatment (Fig. 4C). Within the same timeframe, the primarily freely diffusing BAK1 (Suppl. Fig. S8C) shows an opposing behavior of decreasing density within its clusters (Fig. 4C) as well as more free and less confined diffusion (Suppl. Fig. S8C). The highly mobile cell wall sensor RLP44 shows no reaction to PSK treatment (Fig. 4B-C, Suppl. Fig. S8D). A conceivable mechanism would be that PSK triggers the formation of tighter PSKR1 clusters, wherein interaction with other receptor complex components such as BAK1 can be facilitated due to mutual proximity. BAK1 is probably residing in separate, smaller clusters from where it is released upon PSK stimulus. The increase in free diffusion may increase the likelihood of freely diffusing BAK1 coming into vicinity with a PSKR1 cluster where BAK1 may be recruited (as proposed similarly in von Arx et al., 2026), so that receptor and co-receptor can transphosphorylate and initiate signaling. The interaction of PSKR1 and BAK1 then enables activation of the proton pumps AHA1 and AHA2, which leads to a drop in apoplastic pH enabling severing of cellulose and cellulose-hemicellulose crosslinks. Since PSKR1 also interacts with the calcium channel CNGC17 (Ladwig et al., 2015), it appears highly probable that the pectin network is altered via CNGC17 upon PSK perception. This culminates in a multifaceted alteration of mechanical cell wall properties that includes the Young’s modulus perpendicular to the cell surface which is accessible by AFM. RLP44 has been shown to be an integral part of this pathway, however we found no indications of a nanoscale mechanism for RLP44 during PSK signaling. We assume that RLP44 resides closely to other PSK-signaling components in the PM, but determination of the exact interaction mechanism will require further research. The summary of this proposed model for rapid PSK signaling in etiolated hypocotyls is depicted in Fig. 7 (left).

**Figure 7:**
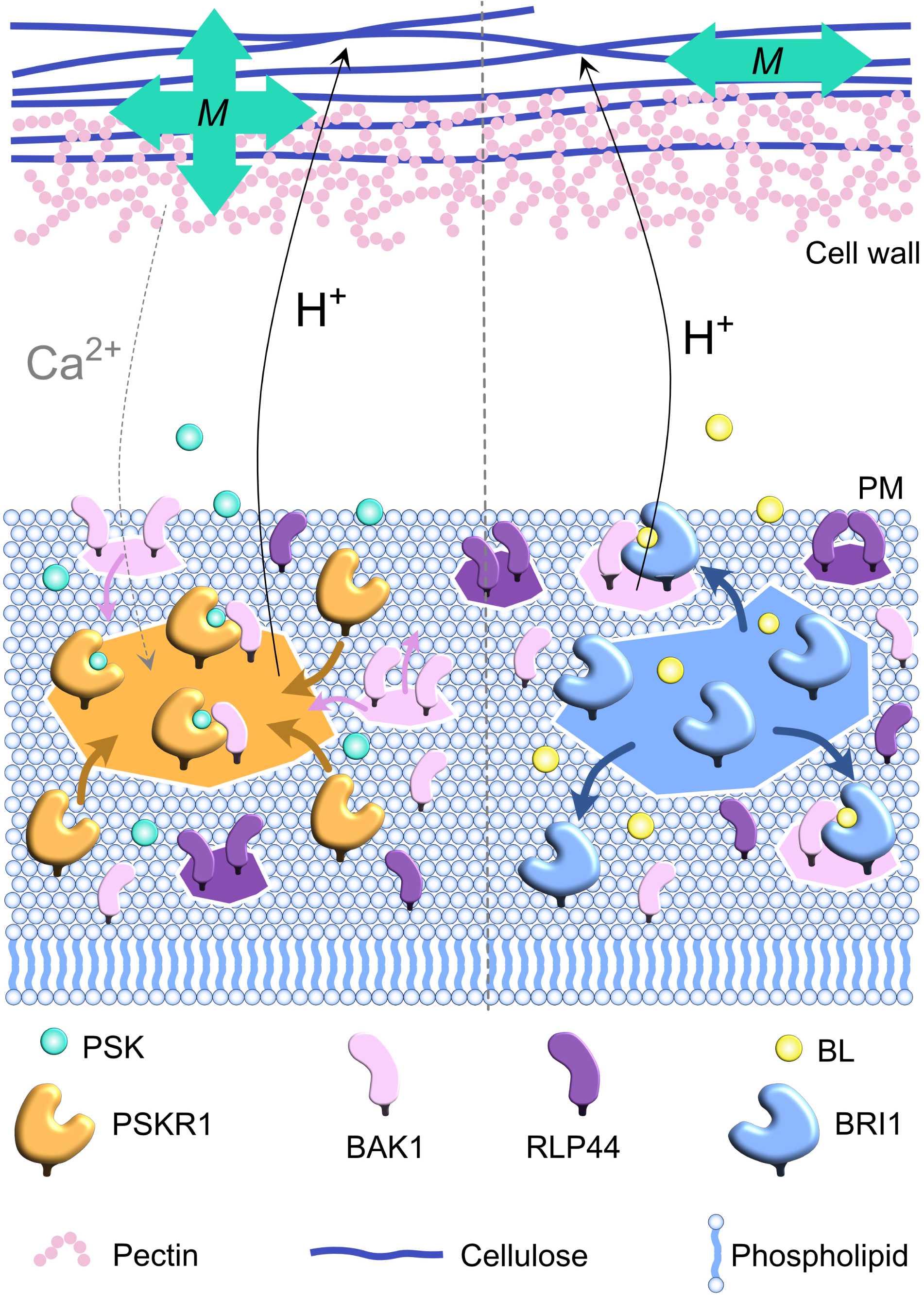
Proposed model of rapid brassinosteroid (BR) and phytosulfokine (PSK) signaling in epidermal cells of etiolated hypocotyls. Left: Schematic representation of rapid PSK signaling in etiolated hypocotyl cells. The ligand PSK is perceived by the PSKR1-BAK1 receptor complex in functional nanoclusters localized in the plasma membrane (PM). PSKR1 receptors relocalize into nanoclusters upon PSK treatment, while BAK1 is released from its initial organization in smaller nanoclusters. The release of BAK1 results in increased free diffusion and less confinement which may aid its availability for temporary recruitment into PSKR1-BAK1 signaling complexes. RLP44 is present in smaller clusters similar to BAK1, which may contribute to PSK signaling on the nanoscale in a non-trivial manner. The perception of PSK induces a release of protons into the apoplast, thereby affecting the cellulose microfibril network within the cell wall. Simultaneously, the pectin network is also affected in an unknown manner that may rely on calcium release. This results in a multifaceted change of mechanical cell wall properties (M). Right: Schematic representation of rapid BR signaling in etiolated hypocotyl cells. BR is perceived by the BRI1-BAK1 receptor complex in functional nanoclusters localized in the PM. Upon BR application, BRI1 receptor cluster density decreases and a second, faster population of BRI1 emerges. Since spatiotemporal organization of BAK1 is oblivious to BR, and BRI1 shows an increase in confined diffusion, BRI1 may assemble into functional receptor complexes by being recruited towards BAK1. RLP44 is present in smaller clusters similar to BAK1, which may contribute to BR signaling on the nanoscale in a non-trivial manner. BR signaling leads to an extrusion of protons into the apoplast, but induces no immediate change within the pectin network. Thereby, the mechanical properties of the cell wall are potentially changed anisotropically. Abbreviations: Brassinosteroid (BR), phytosulfokine (PSK), plasma membrane (PM), mechanical cell wall properties (M).

On the other hand, we found that there is no clear effect on the apparent stiffness measured by AFM after BL treatment (Fig. 1D), despite a receptor complex-dependent lowering of the cell wall pH (Fig. 2C). Since the CT-BDP measurements showed now effect of BL, an effect on the pectin mesh appears unlikely, leaving cellulose as the most probable BR-signaling target within the cell wall. However, since FC resulted in a lowering of apparent stiffness, the mechanism behind BL-induced cell elongation appears to be more complex than general alteration of the cellulose microfibril crosslinks. It should be noted that we performed shallow indentations perpendicular to the cell surface, which probe rather local responses of the cell wall matrix than longitudinal responses like cellulose stretching (Rausch et al., 2026). It is therefore conceivable, that BL induces cell wall changes on an axis unaccessible by our AFM experiments, which may contribute to directional extensibility (see Fig. 7, right). On a molecular level, we found that only the receptor BRI1 exhibits changes within its spatiotemporal organization, unlike BAK1 and RLP44. Importantly, perception of BL triggers a switch within the receptor pool towards a faster moving population, which is accompanied by decreased density within BRI1 clusters and a slight increase in confined behavior. In a scenario that differs from PSK perception, BRI1 may contribute to efficient signaling complex establishment by switching out of its preformed clusters and encountering interaction partners within the smaller BAK1 clusters instead. Faster diffusion may facilitate the encounter with BAK1 clusters, and BRI1 could be temporally recruited towards BAK1. The role of RLP44 within this framework will need to be determined in future research. The summary of our proposed model for rapid BL signaling in etiolated hypocotyls is depicted in Fig. 7 (right).

## Discussion

Cell elongation growth is a fundamental process within plant development that is orchestrated by a multitude of phytohormones. Here we investigated how two closely related signaling complexes contribute to hypocotyl elongation on short timescales. We found a new rapid phenotype linked to PSK signaling using AFM-based indentations of epidermal cells *in vivo* (Fig. 1C). Since the receptor PSKR1 is known to interact with the proton pumps AHA1 and AHA2 similarly as BRI1, as well as the calcium channel CNGC17 (Ladwig et al., 2015), a cell wall-related output of this receptor complex is plausible. PSK signaling has been connected to long-term cell wall effects in previous works. Yu et al. (2016) found that overexpression of the PSK precursor gene AtPSK4 increased the expression of expansin-encoding genes. Zhang et al. (2022) demonstrated how PSK contributes to resistance against *Verticillium dahliae* in cotton by indirect regulation of pectin methylesterification. Our AFM measurements reveal that PSK can affect cell wall properties on short timescales without the involvement of transcriptional responses. Since PSK growth occurs primarly in the epidermis (Hartmann et al., 2013), AFM is a suitable tool to assess PSK-induced cell physiological responses. Furthermore, in the light of recent findings by Krupař et al. (2026) showing that pH is not necessarily an indicator of cell surface properties such as strain, it is worth to assess cellular properties more holistically with multiple independent methods, such as AFM combined with CT-based FLIM.

It is by combining these advanced methods that we found the non-trivial differences between rapid responses to PSK and BL. Even though both treatments lead to a decrease in cell wall pH (Fig. 2C), there was no clear effect of BL observable during AFM indentations. Since we operated our AFM experiments in the shallow regime that mostly probes local variations in the cell wall matrix (Rausch et al., 2026), a distinct PSK-induced alteration of the pectin network appears plausible, as confirmed by our CT-BDP measurements (Fig. 3B). Nevertheless, treatment with the proton pump activator FC also resulted in reduced apparent stiffness in spite of showing no effect on the FLT of CT-BDP (Fig. 3B). The role of pectin in acid growth processes has gained more attention (Arsuffi et al., 2018). For instance, Rui et al. (2017) showed that the dynamic of stomatal opening after FC-treatment is altered when expression of the pectin-degrading POLYGALACTURONASE INVOLVED IN EXPANSION3 is varied. On the other hand, the work by Jones et al. (2003) demonstrated that FC-induced guard cell opening is not inhibited by disturbance of the pectin network. Taken together with our results, the exact mechanism of pectin (and potentially other cell wall components such as hemicellulose) in acid growth processes may be more complex than previously thought.

Another possible explanation for the discrepancy between our treatments examined by AFM may be related to the dimensionality of the rapid BR response. Shallow indentations by AFM (perpendicularly to the cell surface) are not primarily probing wall stretching in the acropetal elongation direction (Rausch et al., 2026). Differences in these properties could, for instance, originate in differential reorientations of microtubules, which serve as movement trajectories for cellulose synthesases (Bringmann et al., 2012; Li et al., 2012). Auxin is known to induce reorientation of microtubules in etiolated hypocotyls from longitudinal to transverse and oblique configurations within 1 h (Adamowski et al., 2019). Longterm exposure to BL differentially affects microtubules in the top and bottom part of etiolated hypocotyls (Somssich et al., 2021). In a recent study by Gallemí et al. (2025), the authors found a decrease of the Young’s Modulus in etiolated seedlings after 2 h treatment with BL as well as FC, however these data were obtained in plasmolyzed seedlings at fairly high concentrations (endogenous levels of most active BRs range from sub-nanomolar to low nanomolar concentrations; Caregnato et al. (2026)). It is therefore conceivable that within the short timeframes investigated in our work here, cellulose is affected in unique ways by BL that will require further research with regard to longitudinal extensibility and microfibril orientation. These properties will require different experimental setups, such as extensometers (Robinson et al., 2017) or deep AFM indentation that probes the longitudinal tension of cellulose fibers as well as turgor pressure and therefore requires mindful data analysis and interpretation (Rausch et al., 2026).

The choice of appropriate, representative concentrations of BL is of specific importance when putting our results into context with previous works. We chose to perform our experiments in the low nanomolar range (10 nM) to represent endogenous levels of BRs (Caregnato et al., 2026) and to represent the regime where the pH-lowering effect of BL reached its maximum in our previous work (Großeholz et al., 2022). We found that the diffusion coefficient of BRI1 increased upon BL treatment, which is in line with the work by Wang et al. (2015b) who observed a similar increase in lateral mobility of GFP-tagged BRI1 in *A. thaliana* roots after treatment with 100 nM BL. It is noteworthy, that Wang et al. (2015b) also observed a two-population distribution with a shift towards higher mobility upon BL stimulus as we observed in our data. However, their imaging of BRI1-GFP particles is not a true single-molecule technique like sptPALM. Similarly, the observation by Bücherl et al. (2017) that the mobility of entire BRI1 clusters is decreased upon BL treatment of cotyledons is based on GFP-tagged protein imaging and therefore not directly comparable with the data presented here. The recent study by von Arx et al. (2026) used sptPALM to evaluate how BRI1 responds to 1 µM BL within 10 min in light-grown hypocotyls of *A. thaliana*. While the authors observed no effect with respect to the nanoscale organization of BRI1 in this setting, the difference in developmental status (light-grown versus dark-grown) as well as the 100-fold difference in ligand concentration complicates a direct comparison with our findings. The findings by von Arx et al. (2026), however, confirm our observation that LRR-RLKs such as BRI1 and PSKR1 have stronger tendencies to form clusters compared to BAK1. Interestingly, von Arx et al. (2026) observed that BAK1 is spatially arrested after both flagellin and BL treatment. While the evaluation methods for the tracking data obtained by sptPALM differ between this work and the publication by von Arx et al. (2026), BAK1 appears to behave differently in dark-grown hypocotyls used in this work. Notably, our treatment with 10 nM BL had no effect on the spatiotemporal patterns of BAK1, but 100 nM PSK triggered a decrease in cluster density and a tendency towards increased free diffusion. This could indicate that spatial arrest and recruitment into nanodomains as proposed by von Arx et al. (2026) is sensitive to the developmental context as well as the specific reaction pathway. It is conceivable that BAK1 might reside in clusters spatially separated from PSKR1 that then release BAK1 available for PSK sensing upon ligand application. The importance of BAK1 for signaling complex organization was underlined by the reversed clustering dynamics of PSKR1 in the *bak1-4* mutant background. The co-receptor may therefore act as a molecular glue recruited into PSKR1-enriched domains, so that interaction, transphosphorylation, and ultimately the induction of cell wall altering signaling processes are enabled.

Currently, the hypothesis of PSKR1 and BAK1 coming together in the same nanodomain can neither be confirmed nor refuted by sptPALM experiments. The current gold standard fluorophore for sptPALM in plants, mEos3.2 (zur Oven-Krockhaus et al., 2026), only allows tracking of one protein at a time, which means that discrimination between interacting and non-interacting receptor complex components is not possible. Dual-color sptPALM, in which photoactivatable fluorophores such as PA-GFP and PATag-RFP allow for simultaneous tracking of two proteins, has been recently introduced (Rohr et al., 2026). But this method is currently limited to the analysis of diffusion coefficients and still requires further development to enable the analysis of potentially co-localized or spatially associated clusters. We could however show that the early events of rapid signaling processes, such as described here for PSK, do rely on specific single-molecule-level reorganization of respective signaling complex components in the PM.

Another signaling component that may offer interesting insights in the future is RLP44. While we found that it is required for rapid processes such as PSK-induced cell wall alterations, we observed no changes in nanoscale motion patterns of RLP44 for neither of our treatments. RLP44 interacts with BRI1 depending on RLP44’s phospho-state (Garnelo Gómez et al., 2021), so it is conceivable that there are two phosphorylation-dependent subpopulations within our datasets that would only be revealed when performing sptPALM tracking for both forms separately. The low clustering tendency resembles the clustering of BAK1. But since we observed that RLP44 was the most mobile protein in terms of diffusion coefficient and displayed by far the highest percentage of free diffusion, it is possible that the nanoscale interaction mechanism with LRR-RLKs like BRI1 and PSKR1 differs from BAK1.

Based on all our findings, we propose a theoretical model for how rapid responses to BL and PSK are carried out in epidermal cells of etiolated hypocotyls (Fig. 7). Specifically, we elucidated distinct nanoscale organizational patterns and their evolution during early signaling events for relevant signaling complex components and found specificity emerged not only in their single-molecule behavior but also in the cell physiological outcomes realized in the cell wall. We thereby introduce new levels of signal coordination within rapidly elongating hypocotyl cells. The corroboration of this model will profit from the further development of advanced super-resolution methods such as dual-color sptPALM. We particularly encourage the incorporation of the nanoscale information obtained from single-molecule studies into computational frameworks for qualitative descriptions of growth, such as Großeholz et al. (2022). By combining cell physiological experimentation, computational modeling and sptPALM, we anticipate the prediction and precise description of new mechanisms contributing to elongation growth.

## Methods

### Genetic constructs and plant transformation

Fusions of BRI1, PSKR1, BAK1 and RLP44, each under their native promotor with the photoconvertible fluorophore mEos3.2, were used in this study. The generation of the genetic constructs for BRI1, PSKR1 and RLP44 via GoldenGate cloning has been described in earlier publications (Rohr et al., 2024b; Rohr et al., 2024a). The BAK1-mEos3.2 construct has been kindly provided by Dr. Birgit Kemmerling (ZMBP, Tübingen). For sptPALM experiments involving PSKR1 and BRI1, transgenic lines were generated by the floral dip method (Zhang et al., 2006). Specifically, the mutant line *bri1-301* was transformed with the BRI1-mEos3.2 construct and the mutant lines *pskr1-3 pskr2-1* and *bak1-4* were transformed with the PSKR1-mEos3.2 construct.

### Plant material and handling

All plants used in this study were created from *Columbia-0* (*Col-0*) accession of *Arabidopsis thaliana*. The mutant lines *pskr1-3 pskr2-1* (Stührwohldt et al., 2011; Kutschmar et al., 2009), *bri1-301* (Lv et al., 2020; Zhang et al., 2018), *bak1-4* (Chinchilla et al., 2007) and *rlp44^cnu2^* (Wolf et al., 2014) have been previously characterized. The plant line expressing *pBAK1-cBAK1-mEos3.2* in the *bak1-4* background was kindly gifted by Dr. Hannah Weber. Since this line is segretating, seeds were tested for expression of the selection marker pFAST (red) before germination. The plant line expressing *pRLP44-cRLP44-mEos3.2* was kindly gifted by Dr. Leander Rohr and has been described previously in Rohr et al. (2024b). Seeds were sterilized in 70 % Ethanol supplied with 0.01 % Triton-X100 for 30 min, followed by 10 min in 100 % Ethanol before drying on filter paper. Sterile seeds were planted on 1/2 Murashige and Skoog (MS) Medium supplemented with 0.8 % Phytoagar and 0.5 g/l 2-(N-Morpholino)-ethane sulphonic acid (MES). MS media were supplemented with 1 % sucrose for AFM and sptPALM experiments, which was omitted for CarboTag FLIM experiments according to previously published works (Besten et al., 2025; Baez et al., 2026). Seeds were stratified at 4 °C for at least 24 h, then exposed to light for 4-6 h to induce germination, followed by vertical growth for 7 days in the dark to induce etiolation.

### Atomic force microscopy (AFM) measurements

To determine the mechanical properties of periclinal cell walls in the plant hypocotyl, seedlings were carefully transferred with tweezers and immobilized on the bottom of a plastic Petri dish by gently placing them on a narrow streak of vacuum grease. Samples were placed so that the central region of the hypocotyl was accessible and not obscured by the grease. The samples were prevented from sample drift and then covered with water containing the appropriate control or hormone treatment. A commercial AFM setup (MFP3D-BIO, Asylum Research, Santa Barbara, CA, USA) mounted on an inverted optical microscope (Ti-S, Nikon, Tokyo, Japan) and cantilevers with 1 µm radius spherical tips and a nominal spring constant of 200 pN/nm (biosphere B1000-CONT, nanotools GmbH, München, Germany) were used for all AFM measurements. The spring constants of the cantilevers were calibrated using the thermal noise method (Cook et al., 2006). Force maps (30×30 µm², 30×30 pixels) were recorded in the central region of the hypocotyl with a retract distance of 2 µm, a force curve rate of 4.8 Hz, and a trigger force of 50 nN (resulting in typical sample indentations of ≈100 nm). An exemplary force map and topological scan are shown in Suppl. Figure S9. Data were analyzed in Igor Pro (WaveMetrics, Lake Oswego, OR USA) using the Hertz model. Force maps typically contained more than one cell. Therefore, regions of interest (ROIs) were drawn manually and the median stiffness value for each ROI was calculated.

### CarboTag synthesis

We synthesized the CarboTag constructs CT-OG and CT-BDP used in this study according to the protocol published in Besten et al. (2025). In brief, the precursor neopentyl glycol-protected pyridine-4-boronic acid is functionalized with a clickable alkyne group, resulting in the structure termed CarboTag. Subsequently, azide-functional OG and BDP were conjugated to the CarboTag via click chemistry (Kolb et al., 2001), as reported in Besten et al. (2025). The products obtained from click reactions were purified using a C-18 reverse-phase column.

### CarboTag FLIM measurements

CarboTag FLIM measurements were performed on 7-day-old etiolated hypocotyls. Samples were incubated in a 10 µM solution of CT-OG or CT-BDP for 1 h. Samples were then gently transferred with tweezers into the treatment solution consisting of either pure DMSO or a solution of BL, PSK or FC in DMSO for 30 min (see Fig. 2A). This second incubation was omitted for the control experiment for CT-BDP experiments. Samples were then mounted onto a drop of DMSO and covered with a coverslip. The second incubation step and mounting was also tested with aqueous solvents, however, all water-based solutions washed the CT out of the samples and prevented successful imaging (data not shown). Confocal imaging and FLIM measurements were conducted with a Zeiss LSM880 confocal microscope (Carl Zeiss Microscopy GmbH, Jena, Germany) using the Zeiss ZEN black and SymPhoTime software (PicoQuant GmbH, Berlin, Germany). Confocal images were acquired using a 40x water-immersion objective (NA = 1.2).

Samples were excited with a 488 nm argon laser at 18 % laser power and fluorescence emission was detected between 500 - 550 nm. Images were recorded at a resolution of 256 x 256 pixels over a 35.42 µm x 35.42 µm field of view.

The PicoQuant FLIM unit used for FLIM measurements is equipped with a PicoQuant LDH-P-C-485B laser, a Sepia PDL 828-L multichannel diode laser driver, a PMA Hybrid 40 detector and a TimeHarp 260 NANO Dual time-correlated single-photon counting (PicoQuant GmbH, Berlin, Germany). FLIM measurements were stopped once the brightest pixel reached a photon count of 200. Subsequent analysis was performed in SymPho-Time 64 (PicoQuant GmbH, Berlin, Germany). The free ROI function was used to select only apoplastic regions of samples. FLT calculation was performed via n-exponential reconvolution for fitting with n=3, using the calculated instrument response function. The average FLT of samples was determined by using one initial fit followed by two additional fits. Further statistical analysis was performed in custom-made RStudio pipelines.

### Single-particle tracking photoactivated localization microscopy (sptPALM)

The super-resolution setup used in this work has been previously described in Rohr et al. (2024b). SptPALM experiments were performed in 7-day-old etiolated *A. thaliana* seedlings expressing the protein of interest tagged with mEos3.2 (see section Genetic constructs and plant transformation Methods). Samples were incubated in liquid MS-media containing BL, PSK or the control treatment for 30 min before mounting. Seedlings were gently placed between two coverslips with a drop of water and then mounted onto the specimen stage with a brass ring to gently flatten the sample. Laser instrumentation and filters used for photoconversion and imaging of mEos3.2 have been previously described in Rohr et al. (2024b).

Samples were positioned in the appropriate focal plane with optimum VAEM angles for single-molecule tracking in a larger 51.2 µm x 15.2 µm observation area with a 10 Hz frame rate, before switching to a smaller 12.8 µm x 12.8 µm observation area for recording. Movies were acquired within the smaller window at a frame rate of 50 Hz with 2500 frames per movie. For each measurement day, a dark-noise calibration file was recorded using identical acquisition settings.

### Analysis of single-molecule tracking data

Data analysis was performed in OneFlowTraX (Rohr et al., 2024b). Files were inspected individually for obvious outliers to be excluded from analysis before being subjected to Batch Analysis (Rohr et al., 2024b). Localization, tracking and MSD analysis was performed according to the parameters for mEos3.2 listed in Rohr et al. (2024b). Files were visually inspected both individually and grouped during mobility analysis to account for multiple populations if adequate. For BRI1, the distinct diffusion coefficients and relative fraction of the two subpopulations were identified via a two-component Gaussian mixture model using the component-fit setting of OneFlowTraX (Rohr et al., 2024b). Cluster size determination was performed using Voronoi tessellation (Levet et al., 2025), with a minimum of 5 tracks per cluster and a relative density threshold *α ≥* 2 . Since cluster areas followed a log-normal distribution, we performed the statistical comparison between datasets according to the method suggested by Zhou et al. (1997) for lognormally distributed data. All further data analysis was performed in custom-made applications in RStudio.

### Motion class analysis

Motion classification was performed according to the ’divide-and-conquer moment scaling spectrum’ (DC-MSS) algorithm introduced by Vega et al. (2018), as previously reported in Rohr et al., 2024a. Trajectories consisting of at least 20 localizations were considered for motion classification by DC-MSS. Therein, tracks are initially segmented into shorter fragments based on maximum pairwise distance within fragments. These segments are then classified by moment scaling spectrum (MSS) analysis into either directed diffusion, free diffusion, confined diffusion or immobility. The MSS analysis algorithm presented in Vega et al. (2018) relies on optimized MSS slope thresholds specifically derived to accurately capture the immobile state (in addition to confined motion).

### Statistical analysis and visualization

Normality was assessed using the Shapiro-Wilk test, and statistical significance between datasets was determined using the Wilcoxon test. Statistical tests and visualization of datasets were performed in R version 4.4.1 R Core Team (2021). Figure 2A was created in BioRender. Rausch, L. (2026) https://BioRender.com/f33tz3a. Figure 7 was created in Affinity Designer v. 2026 3.2.1. Figure assembly was performed in Adobe Illustrator v. 2025 29.3.1 and Affinity Designer v. 2026 3.2.3.

### Data availability

All data are available in the main text or the supplemental information.

## Author contributions

LR: Conceptualization, Data curation, Formal analysis, Investigation, Visualization, Writing – original draft, Writing – review & editing. AB: Data curation, Formal analysis, Investigation, Writing – review & editing. DZ: Data curation, Investigation, Writing – review & editing. RMD: Resources, Writing - review & editing. JS: Resources, Writing - review & editing. SzOK: Resources, Writing - review & editing. HFB: Resources, Supervision, Writing – review & editing. TES: Conceptualization, Funding acquisition, Resources, Supervision, Writing – review & editing. KH: Conceptualization, Funding acquisition, Supervision, Writing – review & editing.

## Acknowledgments

We thank Dr. Leander Rohr and Dr. Gabriella Mosca for fruitful discussions of the manuscript.

## Supplemental information

### Supplemental figures

**Figure S1:**
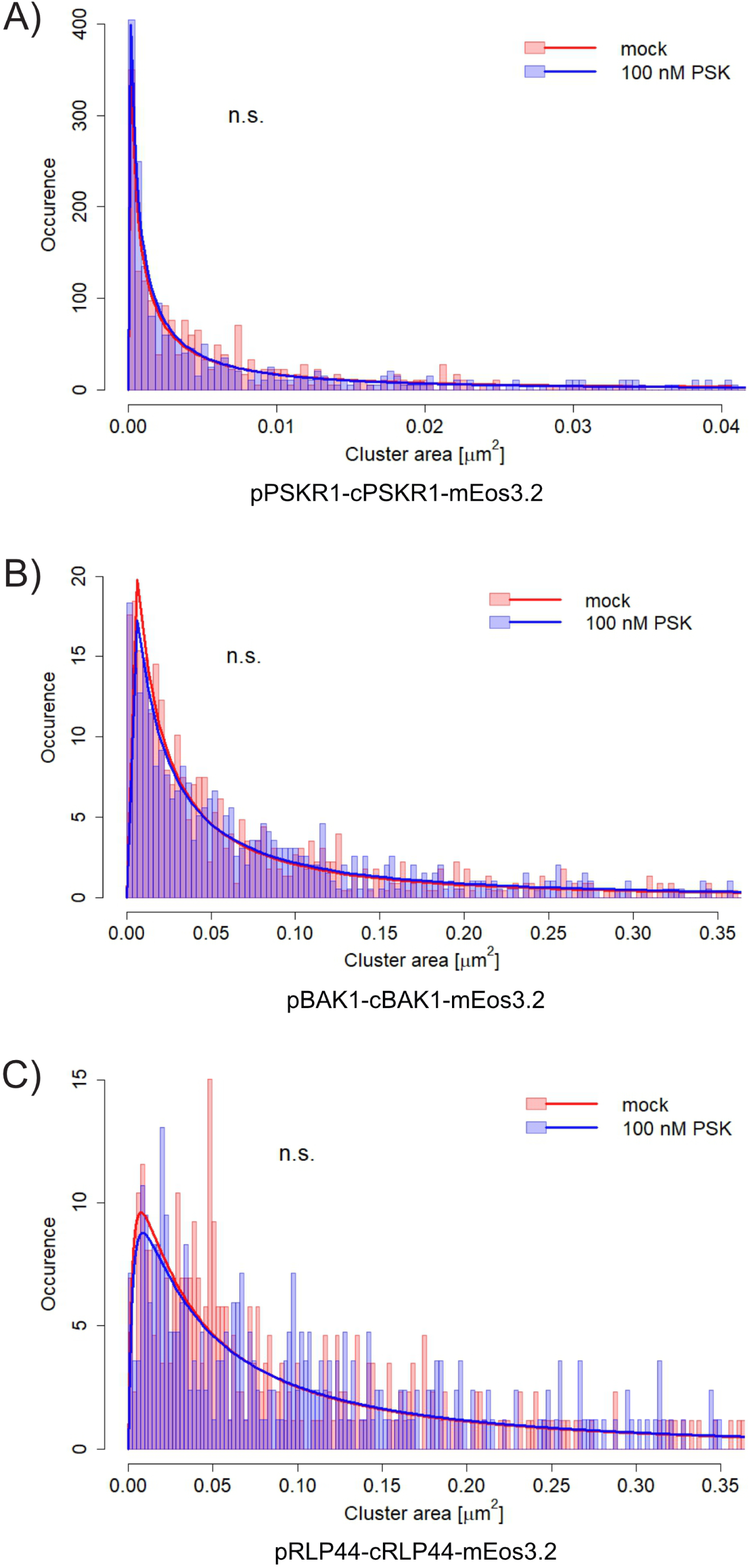
Cluster size distribution (occurrence plotted against area in µm^2^) of phytosulfokine (PSK) signaling components fused to mEos3.2 during signal initiation in epidermal cells of etiolated hypocotyls in *A. thaliana*, related to figure 4. A) Cluster size distribution of PSKR1 after 30 min incubation with 100 nM PSK or control treatment with log-normal fits (pPSKR1-cPSKR1-mEos3.2 in *pskr1-3 pskr2-1*). B) Cluster size distribution of BAK1 after 30 min incubation with 100 nM PSK or control treatment with log-normal fits (pBAK1-cBAK1-mEos3.2 in *bak1-4*). The control data shown here for BAK1 are shared with figure S3B, as both experiments were performed concurrently under identical conditions. C) Cluster size distribution of RLP44 after 30 min incubation with 100 nM PSK or control treatment with log-normal fits (pRLP44-cRLP44-mEos3.2 in *Col-0*). The control data shown here for RLP44 are shared with figure S3C, as both experiments were performed concurrently under identical conditions. All data is pooled from 4 independent measurement days. The number of cells per condition in which cluster areas were evaluated is n *≥* 41 for PSKR1, n *≥* 63 for BAK1 and n *≥* 60 for RLP44. Statistical comparison of log-normal distributions was performed according to Zhou et al. (1997). Abbreviations: Phytosulfokine (PSK).

**Figure S2:**
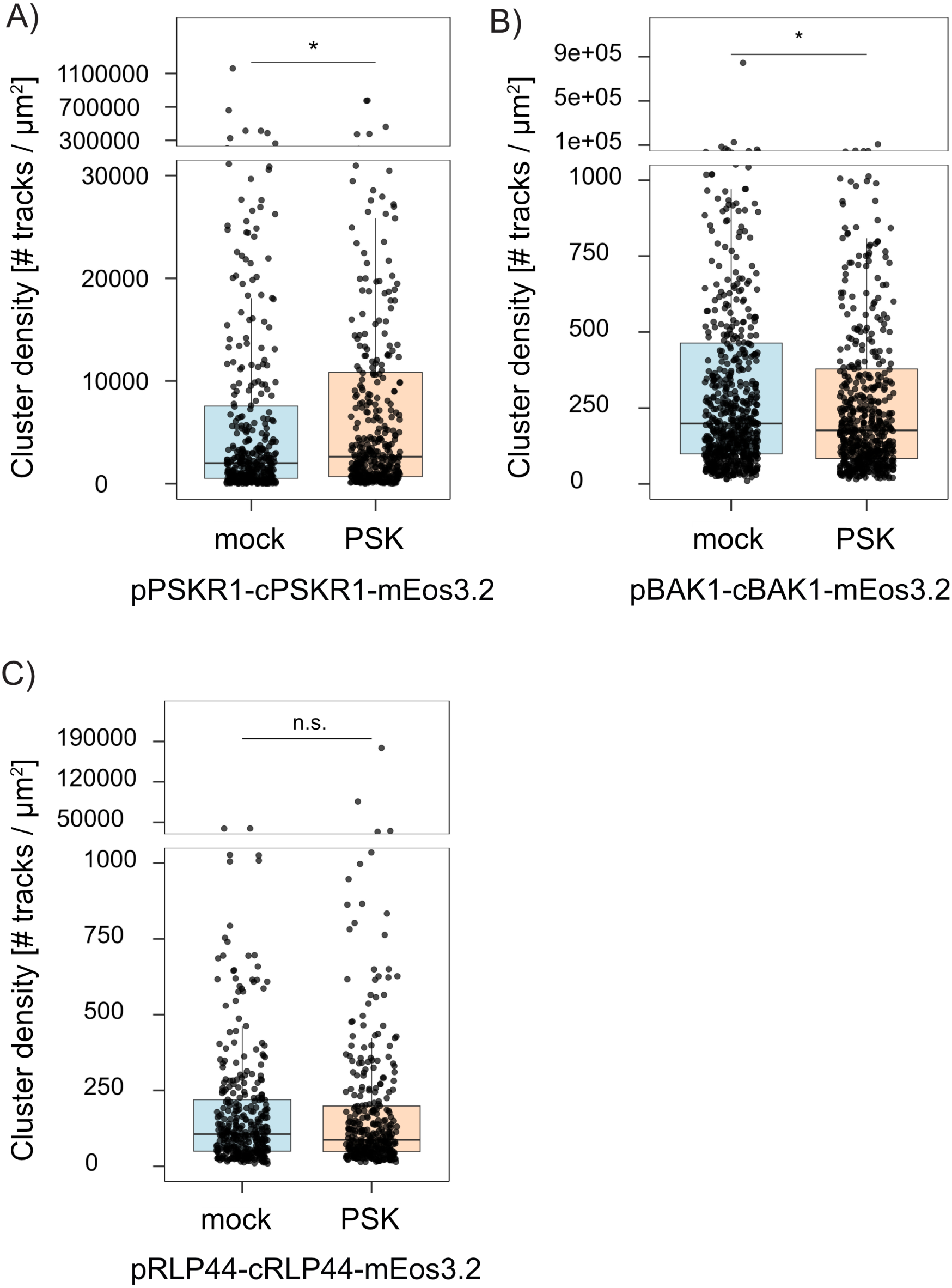
Cluster density distribution of phytosulfokine (PSK) signaling components during signal initiation in epidermal cells of etiolated hypocotyls in *A. thaliana*. Density was calculated as the number of tracks within a cluster divided by its area, related to figure 4. A) Cluster density distribution of PSKR1 after 30 min incubation with 100 nM PSK or control treatment (pPSKR1-cPSKR1-mEos3.2 in *pskr1-3 pskr2-1*). B) Cluster density distribution of BAK1 after 30 min incubation with 100 nM PSK or control treatment (pBAK1-cBAK1-mEos3.2 in *bak1-4*). The control data shown here for BAK1 are shared with figure S4B, as both experiments were performed concurrently under identical conditions. C) Cluster density distribution of RLP44 after 30 min incubation with 100 nM PSK or control treatment (pRLP44-cRLP44-mEos3.2 in *Col-0*). The control data shown here for RLP44 are shared with figure S4B, as both experiments were performed concurrently under identical conditions. All data is pooled from 4 independent measurement days. The number of cells per condition in which cluster densities were evaluated is n *≥* 41 for PSKR1, n *≥* 63 for BAK1 and n *≥* 60 for RLP44. All statistical analyses were performed using custom-made R scripts and the Wilcoxon test distribution (applicability checked prior using a Levene’s test and a Shapiro-Wilk test). p *≤* 0.001 (***); p *≤* 0.01 (**); p *≤* 0.05 (*); p *>* 0.05 (n.s.). Abbreviations: Phytosulfokine (PSK).

**Figure S3:**
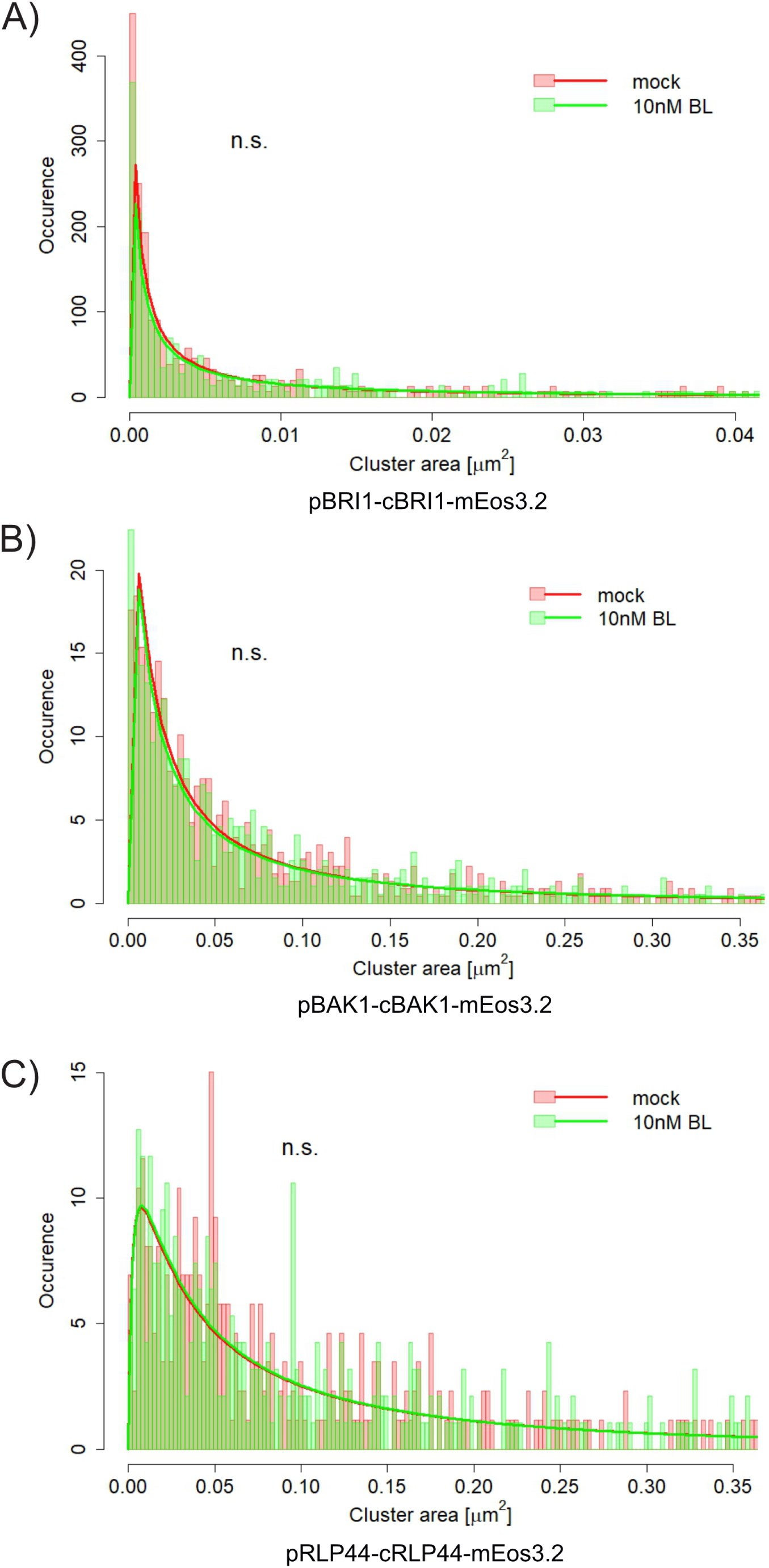
Cluster size distribution (occurrence plotted against area in µm^2^) of brassinosteroid (BR) signaling components fused to mEos3.2 during signal initiation in epidermal cells of etiolated hypocotyls in *A. thaliana*, related to figure 5. A) Cluster size distribution of BRI1 after 30 min incubation with 10 nM brassinolide (BL) or control treatment with log-normal fits (pBRI1-cBRI1-mEos3.2 in *bri1-301*). B) Cluster size distribution of BAK1 after 30 min incubation with 10 nM BL or control treatment with log-normal fits (pBAK1-cBAK1-mEos3.2 in *bak1-4*). Control data for BAK1 are identical to those in figure S1B due to simultaneous execution under identical conditions. C) Cluster size distribution of RLP44 after 30 min incubation with 10 nM BL or control treatment with log-normal fits (pRLP44-cRLP44-mEos3.2 in *Col-0*). Control data for RLP44 are identical to those in figure S1C due to simultaneous execution under identical conditions. All data is pooled from 4 independent measurement days. The number of cells per condition in which cluster areas were evaluated is n *≥* 63 for BRI1, n *≥* 62 for BAK1 and n *≥* 60 for RLP44. Statistical comparison of log-normal distributions was performed according to Zhou et al. (1997). Abbreviations: Brassinosteroid (BR), brassinolide (BL).

**Figure S4:**
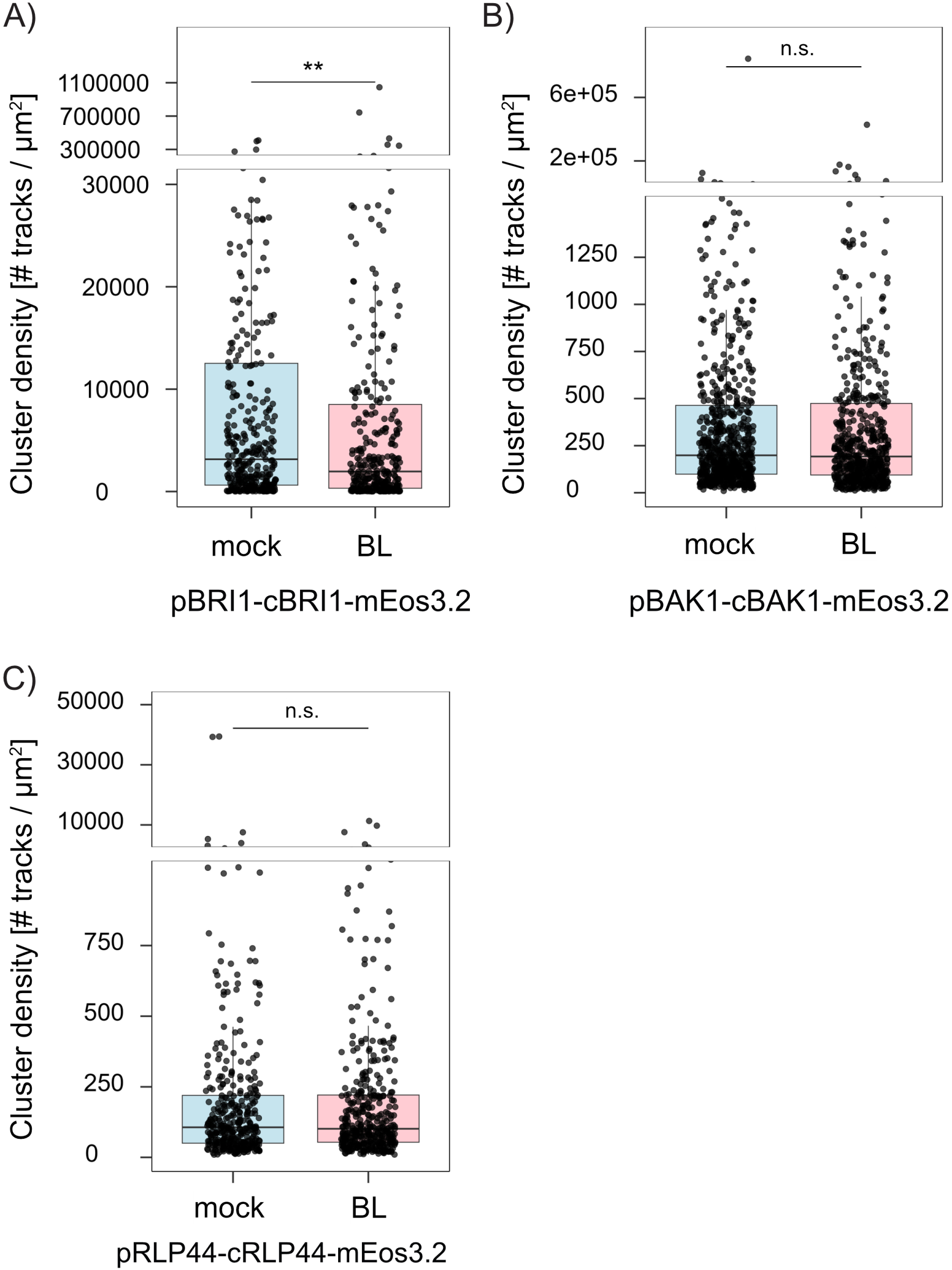
Cluster density distribution of brassinosteroid (BR) signaling components during signal initiation in epidermal cells of etiolated hypocotyls in *A. thaliana*. Density was calculated as the number of tracks within a cluster divided by its area, related to figure 5. A) Cluster density distribution of BRI1 after 30 min incubation with 10 nM brassinolide (BL) or control treatment (pBRI1-cBRI1-mEos3.2 in *bri1-301*). B) Cluster density distribution of BAK1 after 30 min incubation with 10 nM BL or control treatment (pBAK1-cBAK1-mEos3.2 in *bak1-4*). Control data for BAK1 are identical to those in figure S2B due to simultaneous execution under identical conditions. C) Cluster density distribution of RLP44 after 30 min incubation with 10 nM BL or control treatment (pRLP44-cRLP44-mEos3.2 in *Col-0*). Control data for RLP44 are identical to those in figure S2B due to simultaneous execution under identical conditions. All data is pooled from 4 independent measurement days. The number of cells per condition in which cluster densities were evaluated is n *≥* 63 for BRI1, n *≥* 62 for BAK1 and n *≥* 60 for RLP44. All statistical analyses were performed using custom-made R scripts and the Wilcoxon test distribution (applicability checked prior using a Levene’s test and a Shapiro-Wilk test). p *≤* 0.001 (***); p *≤* 0.01 (**); p *≤* 0.05 (*); p *>* 0.05 (n.s.). Abbreviations: Brassinosteroid (BR), brassinolide (BL).

**Figure S5:**
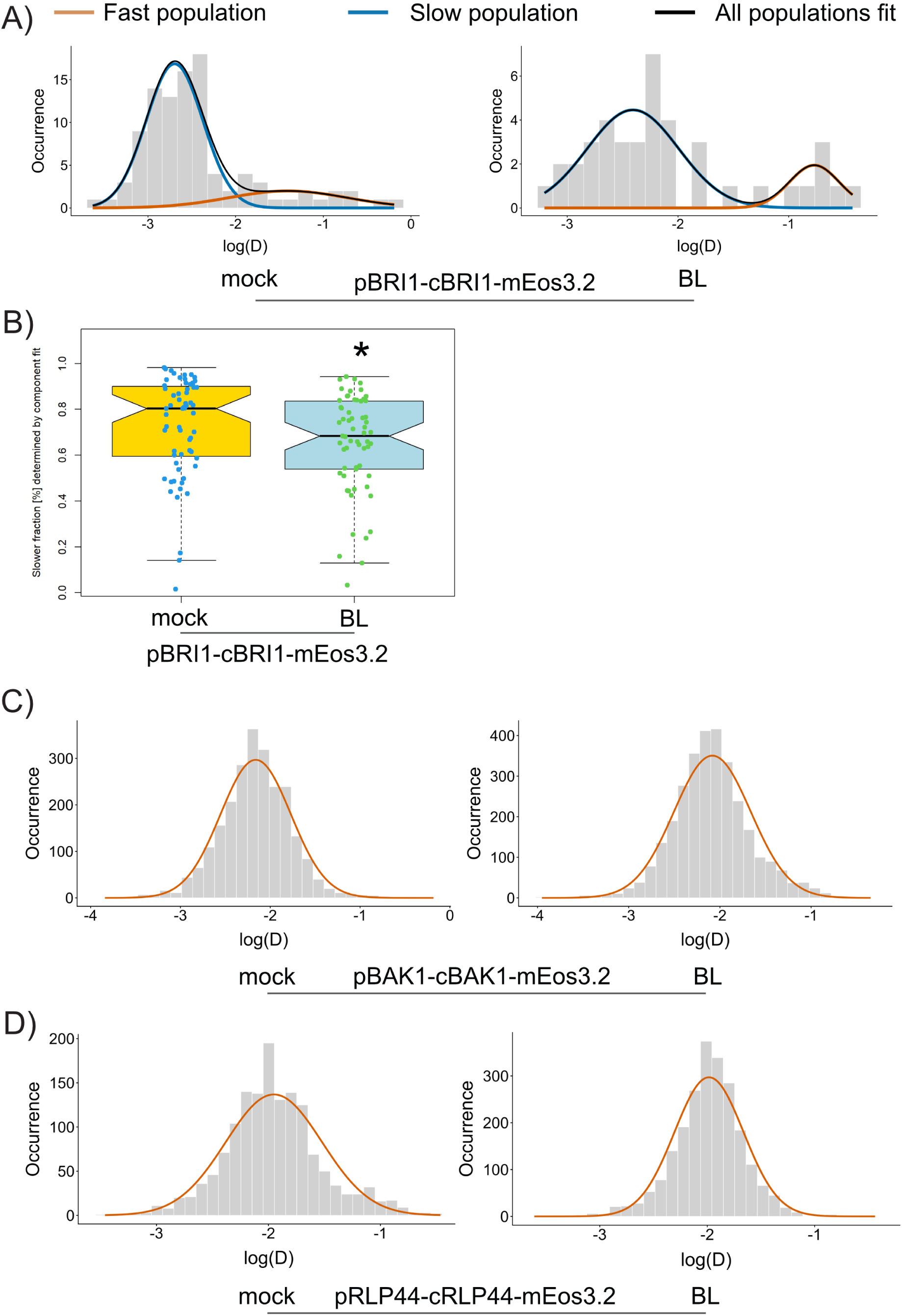
Population analysis of brassinosteroid (BR) signaling components during signal initiation in epidermal cells of etiolated hypocotyls in *A. thaliana*, related to figure 5. A) Exemplary diffusion coefficient distribution of BRI1-mEos3.2 after 30 min incubation with 10 nM brassinolide (BL) or control treatment in individual tracking files obtained from individual cells (pBRI1-cBRI1-mEos3.2 in *bri1-301*). The two populations are indicated by the colored curves obtained via a two-component Gaussian mixture model. B) Fraction of tracks of BRI1-mEos3.2 in slower fraction after 30 min incubation with 10 nM BL or control treatment (determined using the component-fit option in OneFlowTraX (Rohr et al., 2024b)). n = *≥* 63 per condition. C) Diffusion coefficient distribution of BAK1 after 30 min incubation with 10 nM BL or control treatment (pBAK1-cBAK1-mEos3.2 in *bak1-4*, n *≥* 16 cells per condition) with Gaussian fits indicated by orange curves. The control data shown here for BAK1 are shared with figure S6B, as both experiments were performed concurrently under identical conditions. D) Diffusion coefficient distribution of RLP44 after 30 min incubation with 10 nM BL or control treatment (pRLP44-cRLP44-mEos3.2 in *Col-0*, n *≥* 15 cells per condition) with Gaussian fits indicated by orange curves. The control data shown here for BAK1 are shared with figure S6C, as both experiments were performed concurrently under identical conditions. The histograms shown in Fig. S5C-D were obtained from the pooled data for each condition on one exemplary measurement day. Diffusion coefficient histograms were created using custom-made R scripts. Statistical analysis of the data presented in figure S5B was performed using custom-made R scripts and the Wilcoxon test distribution (applicability checked prior using a Levene’s test and a Shapiro-Wilk test). p *≤* 0.001 (***); p *≤* 0.01 (**); p *≤* 0.05 (*); p *>* 0.05 (n.s.). Abbreviations: Brassinosteroid (BR), brassinolide (BL).

**Figure S6:**
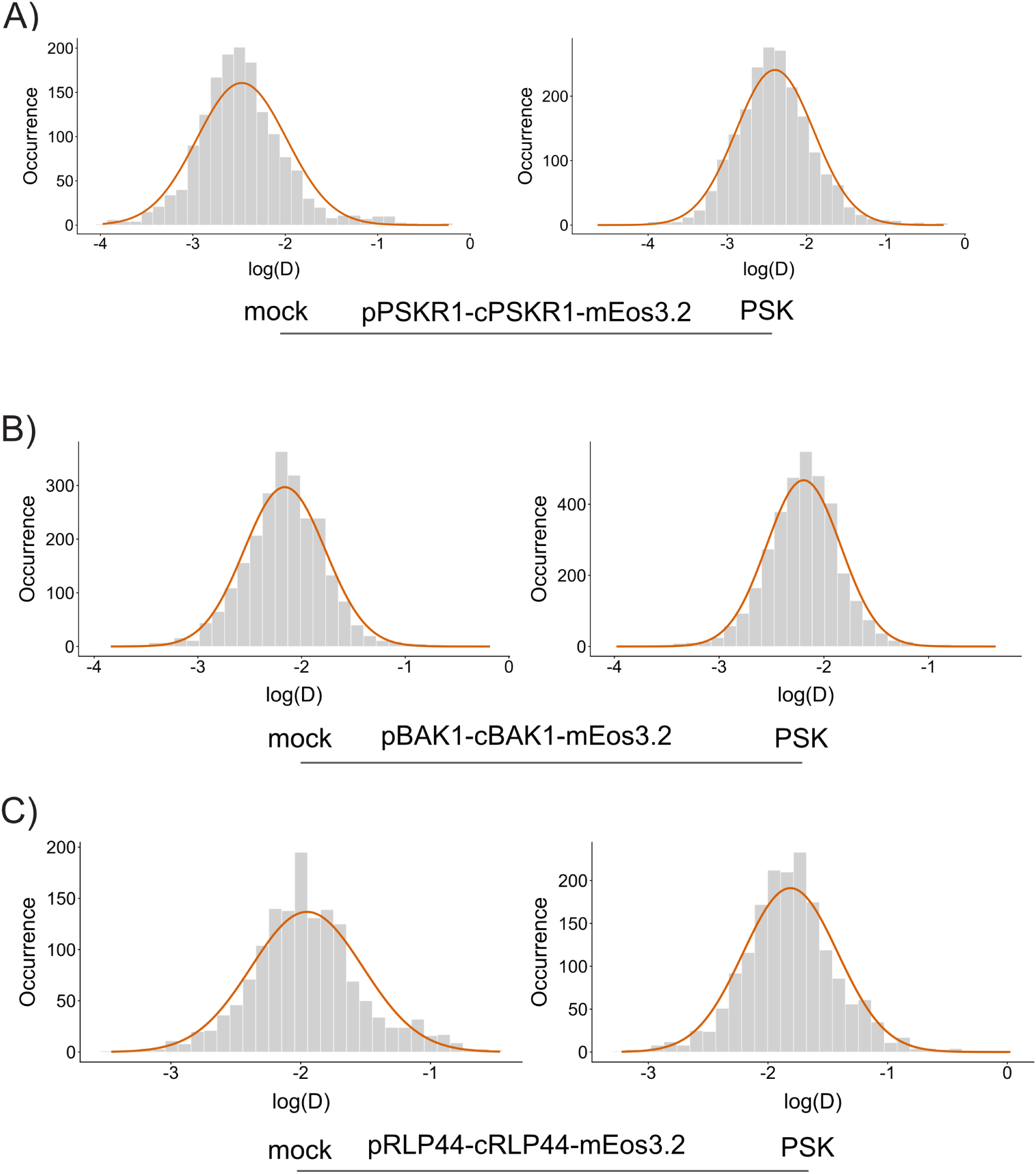
Population analysis of phytosulfokine (PSK) signaling components during signal initiation in epidermal cells of etiolated hypocotyls in *A. thaliana*, related to figure 4. Diffusion coefficient histograms created within the tracking tab of OneFlowTraX (Rohr et al., 2024b) A) Diffusion coefficient distribution of PSKR1 after 30 min incubation with 100 nM PSK or control treatment (pPSKR1-cPSKR1-mEos3.2 in *pskr1-3 pskr2-1*, n *≥* 11 cells per condition) with Gaussian fits indicated by orange curves. B) Diffusion coefficient distribution of BAK1 after 30 min incubation with 100 nM PSK or control treatment (pBAK1-cBAK1-mEos3.2 in *bak1-4*, n *≥* 16 cells per condition) with Gaussian fits indicated by orange curves. Control data for BAK1 are identical to those in figure S5C due to simultaneous execution under identical conditions. C) Diffusion coefficient distribution of RLP44 after 30 min incubation with 100 nM PSK or control treatment (pRLP44-cRLP44-mEos3.2 in *Col-0*, n *≥* 15 cells per condition) with Gaussian fits indicated by orange curves. Control data for RLP44 are identical to those in figure S5D due to simultaneous execution under identical conditions. The histograms shown in Fig. S6A-C were obtained from the pooled data for each condition on one exemplary measurement day. Diffusion coefficient histograms were created using custom-made R scripts. Abbreviations: Phytosulfokine (PSK).

**Figure S7:**
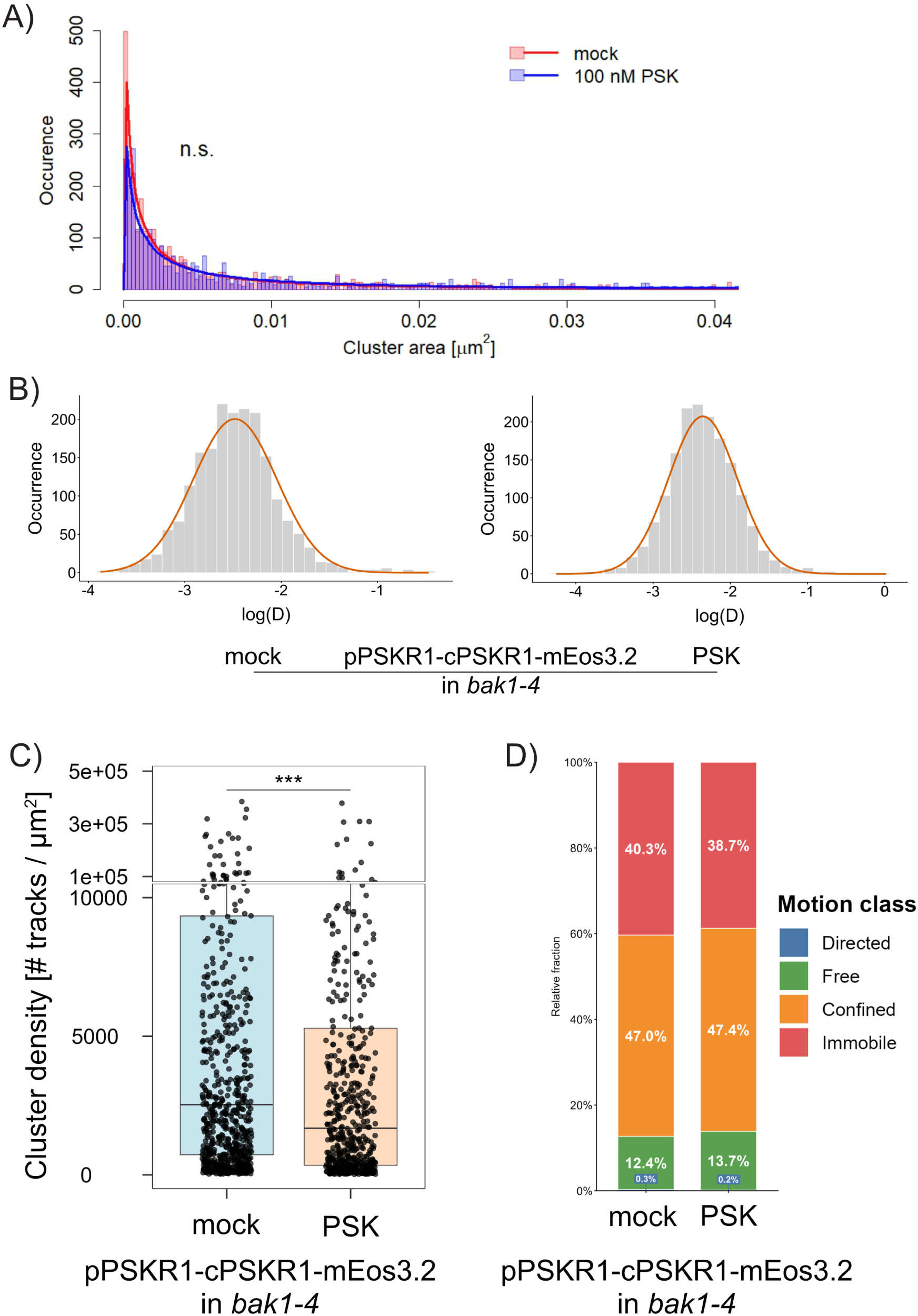
Supplemental data nanoscale spatiotemporal behavior of PSKR1 during Phytosulfokine (PSK) treatment in epidermal cells of 7-day-old etiolated hypocotyls in the mutant background *bak1-4*, related to figure 6. A) Cluster size distribution of PSKR1 after 30 min incubation with 100 nM PSK or control treatment with log-normal fits (pPSKR1-cPSKR1-mEos3.2 in *bak1-4*). B) Diffusion coefficient distribution of PSKR1 after 30 min incubation with 100 nM PSK or control treatment (pPSKR1-cPSKR1-mEos3.2 in *bak1-4*, n *≥* 15 cells per condition) with Gaussian fits indicated by orange curves. C) Cluster density distribution of PSKR1 after 30 min incubation with 100 nM PSK or control treatment (pPSKR1-cPSKR1-mEos3.2 in *bak1-4*). All data is pooled from 4 independent measurement days. Statistical comparison of log-normal distributions shown in figure S7A was performed according to Zhou et al. (1997). D) Proportion of different motion behaviors of PSKR1-mEos3.2 after 30 min incubation with 100 nM PSK or control treatment in epidermal cells of etiolated hypocotyls in *A. thaliana* (pPSKR1-cPSKR1-mEos3.2 in *bak1-4*). Classification of motion types was performed according to Vega et al. (2018). The histograms shown in Fig. S7B were obtained from the pooled data for each condition on one exemplary measurement day. All other data is pooled from 4 independent measurement days. The number of cells per condition in which cluster area in S7A, cluster densities in S7C and motion type classification in S7D were evaluated is n = 60. Diffusion coefficient histograms were created using custom-made R scripts. Statistical analysis of the data presented in figure S7C was performed using custom-made R scripts and the Wilcoxon test distribution (applicability checked prior using a Levene’s test and a Shapiro-Wilk test). p *≤* 0.001 (***); p *≤* 0.01 (**); p *≤* 0.05 (*); p *>* 0.05 (n.s.). Abbreviations: Phytosulfokine (PSK).

**Figure S8:**
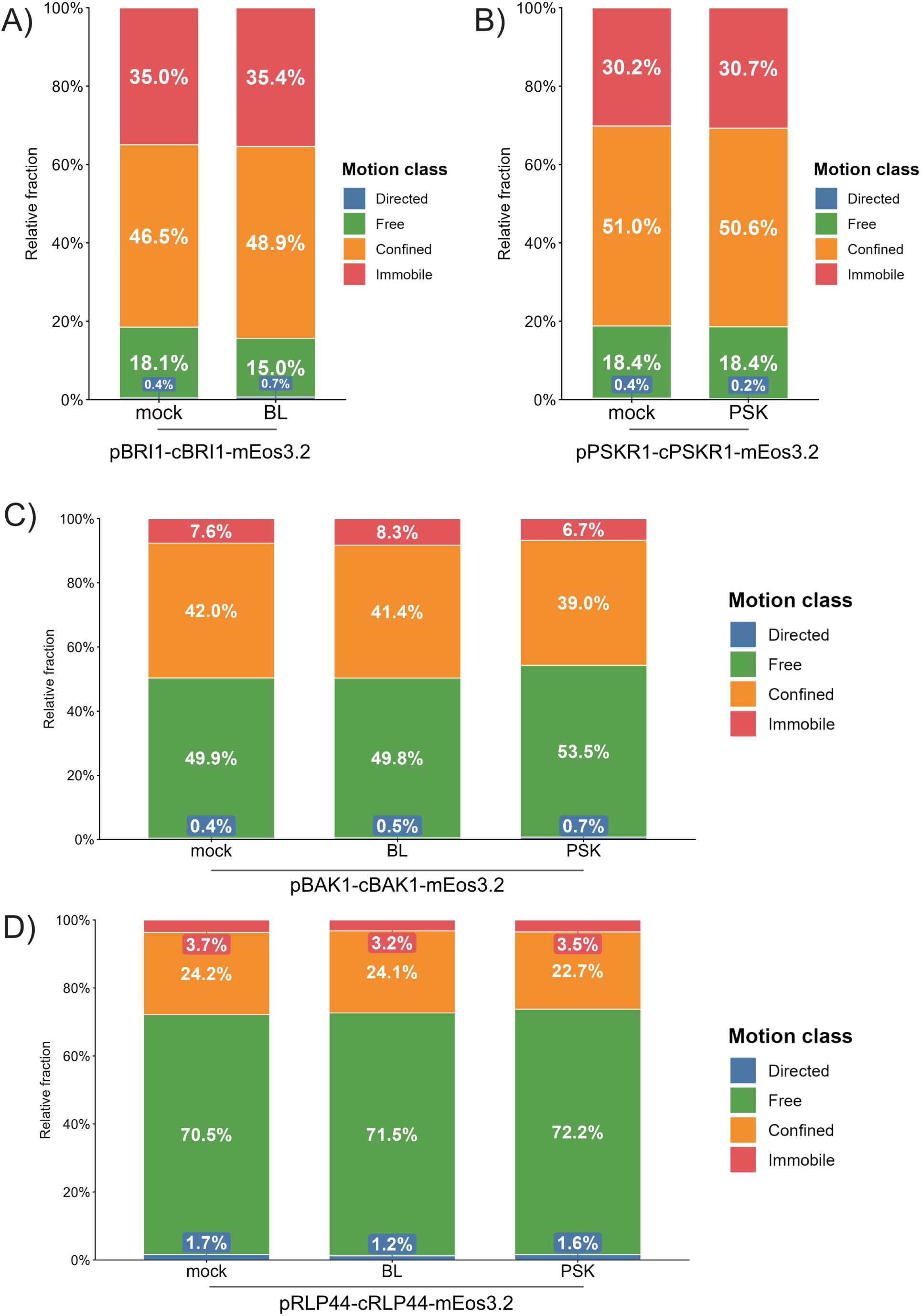
Motion classification of signaling components fused to mEos3.2 during signal initiation in epidermal cells of etiolated hypocotyls in *A. thaliana*. Classification of motion types was performed according to Vega et al. (2018) A) Proportion of different motion behaviors of BRI1-mEos3.2 after 30 min incubation with 10 nM brassinolide (BL) or control treatment (pBRI1-cBRI1-mEos3.2 in *bri1-301*) B) Proportion of different motion behaviors of PSKR1 with phytosulfokine (PSK) or control treatment C) Proportion of different motion behaviors of BAK1 after 30 min incubation with 10 nM BL, 100 nM PSK or control treatment D) Proportion of different motion behaviors of RLP44 after 30 min incubation with 10 nM BL, 100 nM PSK or control treatment. All data is pooled from 4 independent measurement days. The number of cells per condition in which motion classification of tracks was performed is n *≥* 63 for BRI1, n *≥* 41 for PSKR1, n *≥* 62 for BAK1 and n *≥* 60 for RLP44. Abbreviations: Brassinolide (BL), phytosulfokine (PSK).

**Figure S9:**
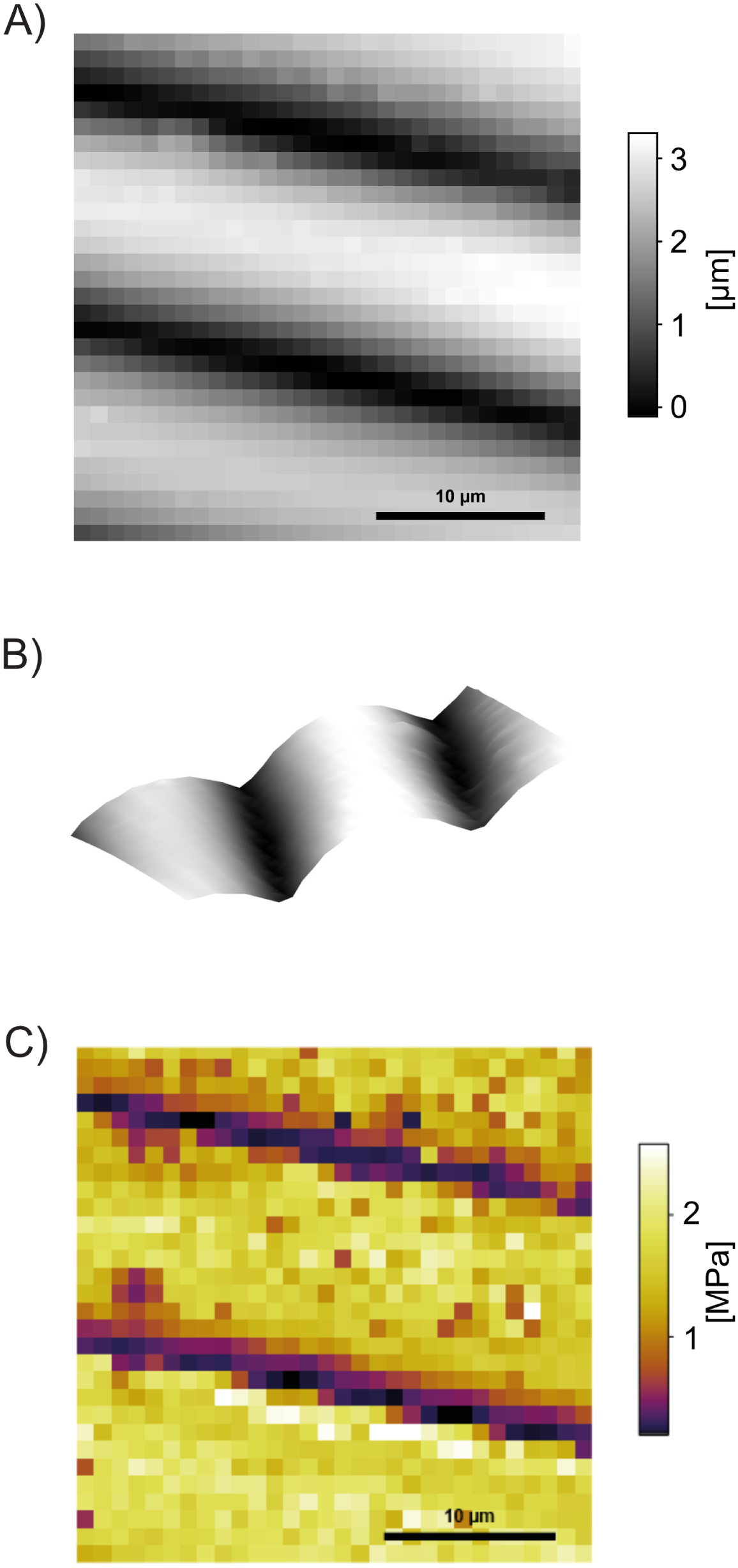
Methodological supplements of atomic force microscopy (AFM) measurements performed on periclinal cell walls in epidermal cells of etiolated hypocotyls in *A. thaliana*. Scans were conducted with a retract distance of 2 µm, a force curve rate of 4.8 Hz, and a trigger force of 50 nN (resulting in typical sample indentations of ≈100 nm). A) Exemplary AFM-derived topological map (30×30 µm², 30×30 pixels) of epidermal cells in etiolated hypocotyls. Scalebar = 10 µm, surface height in µm indicated by grey scale. The shape of the long, ≈10 µm-wide, elongated cells is clearly visible in the topological scan. B) Three-dimensional rendering of the topology of epidermal sample shown in Figure S9A. C) Exemplary AFM-derived force map (30×30 µm², 30×30 pixels). Scalebar = 10 µm, magnitude of apparent stiffness in MPa indicated by color scale. Individual cells were analyzed with regard to apparent stiffness by drawing a region of interest (ROI) manually in the central region of a single cell during analysis in Igor Pro (WaveMetrics, Lake Oswego, OR USA). Then, the median apparent stiffness for each ROI was calculated. Abbreviations: Atomic force microscopy (AFM), region of interest (ROI).

